# The dual actions of the host miRNA-16a in restricting Bovine coronavirus (BCoV) replication through targeting host cell Furin and in enhancing the host immune response

**DOI:** 10.1101/2024.05.29.596392

**Authors:** Abid Ullah Shah, Maged Gomaa Hemida

## Abstract

The roles of host cell miRNAs have not been studied well in the context of BCoV replication and immune regulation. The main aim of this study was to identify some miRNA candidates that regulate essential host genes involved in BCoV replication, tissue tropism, and immune regulation. To achieve these goals, we used two isolates of BCoV (enteric and respiratory) to infect the bovine endothelial cells (BEC) and Madine Darby Bovine Kidney (MDBK) cells. This is in addition to the ex vivo model using the peripheral bovine blood mononuclear cells (PBMC). We determined the miRNA expression profiles in these cells after BCoV infection. miRA-16a is one of the differentially altered during BCoV infection. Our data shows that miRNA-16a is a significantly downregulated miRNA in both in vitro and *ex vivo* models. We confirmed the miRNA-16a expression profile by the qRT-PCR. Overexpression of the pre-miRNA-16a in BEC and MDBK cell lines resulted in marked inhibition of BCoV infection based on the viral genome copy numbers measured by qRT-PCR, the viral protein expression (S and N) measured by Western blot, and the virus infectivity using plaque assay. Our bioinformatic prediction showed that Furin is a potential target for the miRNA-16a. We checked the Furin protein expression level in the pre-miRNA-16a transfected/BCoV infected cells compared to the pre-miRNA scrambled to validate that. Our data shows marked inhibition of the Furin expression levels on the mRNA levels by qRT-PCR and the protein level by Western blot. The BCoV-S protein expression was markedly inhibited on both the mRNA and protein levels. To further confirm the impacts of the downregulation of the Furin enzyme on the replication of BCoV, we used transfected cells with specific Furin-siRNA parallel to the scrambled siRNA. A marked inhibition of BCoV replication was observed in the Furin-siRNA-treated group. To further validate Furin as a novel target for miRNA-16a, we cloned the 3’UTR of the bovine Furin carrying the seed region of the miRNA-16a in the dual luciferase vector. Our data shows luciferase activity in the pre-miRNA-16a transfected cells decreased by more than 50% compared to the cells transfected with the construct carrying the mutated Furin seed region. Our data confirms miRNA-16a inhibits BCoV replication by targeting the host cell Furin and the BCoV-S glycoprotein. It will also enhance the host immune response, which contributes to the inhibition of viral replication. To our knowledge, this is the first study to confirm that Furin is a valid target for the miRNA-16a. Our findings highlight the clinical applications of the host miRNA-16a as a potential miRNA-based vaccine/antiviral therapy.

## 2. Introduction

Bovine coronavirus (BCoV) was recently classified under the order *Nidovirales* in the family *Coronaviridae, the subfamily Orthocoronavirinae, the genus Betacoronavirus,* and the subgenus Embecovirus (1). Both BCoV and the severe acute respiratory syndrome coronaviruses (SARS-CoV)- 1 and 2 belong to the genus Betacoronaviruses. Those viruses share some common characteristics on the phenotypic and genotypic levels. Although BCoV was reported several decades ago, many aspects of viral replication and virus/host interaction have not been explored yet. This contrasts with the SARS-CoV-2, the cause of the COVID-19 pandemic; intensive studies have been carried out and revealed many novel aspects of the SARS-CoV-2/host interaction. Virus entry is a crucial step in the coronavirus replication cycle. The angiotensinogen converting enzyme-2 (ACE2) has been proven to be the functional receptor for SARS-CoV-2 (2). The transmembrane protease serine 2 (TMPRSS2) plays essential roles in the SARS-CoV-2 attachment and entry into the host cells through the cleavage of the viral spike glycoprotein at specific sites. The S protein cleavage activated BCoV infection in the host cell and contributed to virus entry to the host cells (3). The roles of the Neuropilin-1 (NRP-1) in the SARS-CoV-2 entry to the cell have been recently studied (4). The NRP-1 binds to the cleaved substrate of the host cell Furin, which enhances the virus replication and plays an essential role in viral immune evasion (4). However, the roles of Furin, ACE2, TMPRRS2, and NRP-1 in the BCoV replication have not been studied yet. The BCoV genome is a single molecule of positive sense RNA (ssRNA, +Ve). The BCoV genome has the typical genome structure and organization of most coronaviruses. The genome is flanked by two untranslated regions at the 5’ and 3’ ends, respectively. The 5’ two-thirds of the genome consists of a large gene called gene-1, composed of two overlapping open reading frames (ORFs) with a ribosomal frameshift. Gene-1 of BCoV is further processed into 16 non-structural proteins (NSP1-16). The 3’ one-third of the BCoV genome is occupied by five major structural proteins (hemagglutinin esterase (HE), Spike glycoprotein (S), the envelope (E), and the nucleocapsid protein (N)) interspersed with some other non-structural proteins (5). BCoV infection causes several clinical syndromes in the affected cattle, including calf diarrhea and winter dysentery, and it also contributes to the development of the bovine respiratory disease complex, along with other bacterial pathogens (6). Most coronaviruses require initial cleavage steps by some host cell proteases, particularly serine proteases, to initiate an active infection in the target host. Furin cleavage of the SARS-CoV-2-spike is a pre-request for the viral infection in the target hosts (7). Viral tropism heavily depends on the availability of some specific cellular receptors that help the virus enter the host cells and hijack the cellular machinery to favor viral protein synthesis instead of cellular proteins (8). Some other factors may also contribute to this tissue tropism, such as the presence of some auxiliary receptors and the presence of some transcription and translation factors (9). BCoV possesses a dual tissue tropism in the affected cattle (enteric and respiratory) (10). The mechanisms that fine-tune this dual tissue tropism have not been well studied. Host cell microRNAs (miRNAs) are small RNA. molecules (21-25 nucleotides in length) that play important roles in gene regulation at translation levels. The miRNA candidate usually binds to certain regions in the 3’UTR of the target gene, leading to its translation inhibition or repression (11). There is an important region in the structure of each miRNA called the seed region, which is usually located at the positions 2-8 nucleotides at the 5’ end of the miRNA (12). The mechanism of action of each miRNA molecule is mainly governed by the degree of complementarity between the miRNA seed region and the complementary region in each mRNA (13). Some D.N.A. viruses, especially herpes viruses, encode some viral miRNAs (14). In contrast, RNA viruses do not usually end their own miRNAs. However, host cell miRNAs play important roles in both D.N.A. and RNA viral replication, tropism, and immune regulation/evasion (8). Little is known about the role of host cell miRNAs in the molecular biology of BCoV. We recently identified some potential host cell miRNAs that may play essential roles in the molecular pathogenesis of BCoV and could partially explain this virus’s dual tropism phenomenon (15). It was shown that miRNA-16a involves many processes, including cell cycle and tumor formation. The miRNA-16 also acts as a diagnostic marker, is involved in gene regulation, and acts as a potential therapeutic target for hepatitis C virus (HCV) infection in humans (16, 17). However, the roles of miRNA-16a in BCoV replication immune regulation/evasion have not been studied yet. Our data shows that miRNA-16a expression is differentially altered during BCoV/Ent/Resp isolates infection. The main goals of the current study were to confirm the differential expression of miRNA-16a in the context of BCoV infection, to study the impacts of miRNA-16a overexpression on the BCoV replication, and to study the mechanism of action of miRNA-16a in fine-tuning of BCoV replication and immune regulation.

## 3. Materials and Methods

### 3.1. Viruses and Cell Lines

Bovine pulmonary artery endothelial cells (BEC.: ATCC® CRL-1733™) were obtained from ATCC. The BEC. cells were tested for the absence of BVDV. The BEC. cells were cultures in F12 media (ATCC, 30-2004), supplemented with 10% horse serum (H.S.) (Gibco; Ref. No. 26050-088) and 1% 10,000 ug/mL streptomycin and 10,000 units/mL penicillin antibiotics (Gibco; Ref. No. 15140-122). The Madine Darby Bovine Kidney (MDBK) cells were kindly provided by Dr. Udeni B. R. Balasuriya, Louisiana State University). The MDBK cells were cultured in Minimum Essential Medium Eagle media (Sigma-Aldrich, Cat. No. M0200-500ML), supplemented with 10% horse serum and 1% streptomycin and penicillin antibiotics. Cells were incubated at 37°C at 5% CO_2_ for subsequent culture. The Human Embryonic Kidney Cells 293 (HEK-293) were obtained from the ATCC (Catalog # CRL-3216 ™) and used in the dual Luciferase assay as described below (section 3.10). Two bovine coronavirus (BCoV) isolates were used; the enteric isolate ’Mebus’ (18) was obtained from B.E.I. resources (B.E.I. Resources, NIAID, N.I.H.: Bovine Coronavirus (BCoV), Mebus, NR-445). The respiratory isolate of BCoV was kindly provided by Dr. Aspen Workman (Animal Health Genomics Research Unit, USDA, A.R.S., U.S. Meat Animal Research) (19).

### 3.2. The Next Generation Sequencing (NGS) and the host cell miRNA expression profiles during the BCoV replication

The BEC cells were infected independently with either BCoV enteric or BCoV respiratory isolates using a multiplicity of infections (MOI=1). The infected cells were observed under the inverted microscope daily for up to 4 days post-infection (4dpi). We monitored the infected cells for the development of any cytopathic effects (C.P.E.) for up to five days post-infection (5 dpi). Compared to the sham (phosphate buffer saline; PBS) infected cells, cells infected with BCoVs showed some morphological changes, such as rounding up and detachment from the confluent monolayer sheet.

We collected the cell culture supernatants from all groups of infected cells, including the sham infected. We extracted the total miRNAs using the miRNeasy Micro Kit (Qiagen: Cat. No. 217084) per the manufacturer’s instructions. The total miRNAs were submitted to L.C. Sciences (L.C. Sciences, L.L.C., 2575 West Belfort Street, Houston, TX 77054, USA) for the reporting of the miRNA expression profiles in various treated groups of cells. Briefly, all extracted RNA was used in the library preparation following Illumina’s TruSeq -small-RNA-sample preparation protocols (Illumina, San Diego, CA, U.S.A.). The D.N.A. library’s quality control analysis and quantification were performed using Agilent Technologies 2100 Bioanalyzer High Sensitivity D.N.A. Chip. Single-end sequencing 50bp was performed on Illumina’s Hiseq 2500 sequencing system following the manufacturer’s recommended protocols. Differential expression of miRNAs based on normalized deep-sequencing counts was analyzed using various statistical tests, including the Fisher exact test, Chi-squared 2X2 test, Chi-squared nXn test, the Student’s t-test, or ANOVA, depending on the experimental design. We used the Protein Analysis Through Evolutionary Relationships (PANTHER) classification version 9.0 to describe the function and properties of host genes and their related pathways with differential expression. The Kyoto Encyclopedia of Genes and Genomes (KEGG) pathways and heatmaps used in this study were designed using the online data analysis and visualization platform (https://www.bioinformatics.com.cn/en).

### 3.3. Determination of the miRNA expression profiles following BCoV/Ent or BCoV/Rep isolates infection

We tested the MDBK, the BEC, and the PBMC to confirm the miRNA expression profiles obtained from the NGS data. We infected each group of cells with either the BCoV/Ent or BCoV/Resp isolates parallel to the sham (PBS)-infected cells. Following manufacturer instructions, the total miRNAs were isolated using the Pure Link miRNA isolation kit (Invitrogen; REF: K157001). We used the Nanodrop OneC (Thermo Fisher Scientific) to check the quality and concentration of the extracted total RNAs from all treated groups of cells. The complementary D.N.A. (cDNA) and the quantitative Reverse Transcriptase-Polymerase Chain Reaction (qRT-PCR) were performed using All-in-One miRNA qRT-PCR Detection Kit 2.0 (GeneCopoeia; Cat. No. QP115) following manufacturer instructions. All the primers for miRNAs were designed using sRNAPrimerDB (20). The relative expression of miRNAs was normalized to the endogenous reference U6 according to 2^-^^Ct^ method (21). All the miRNA primers used in this study are listed in Table 1.

**Table 1:**
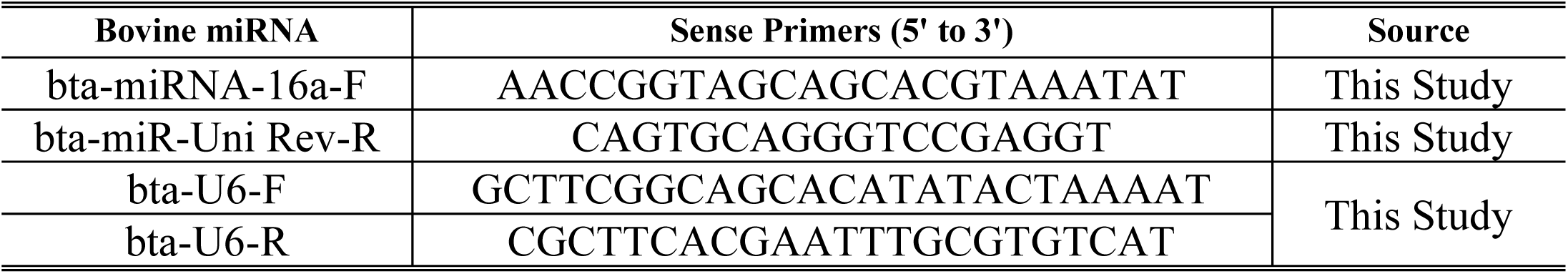
List of the oligonucleotides used for the amplification of miRNA expression profiles.

### 3.4. Host mRNA Extraction, amplification, and Quantification

The total RNAs were isolated from MDBK, BEC, and PBMC and infected and from the control group of cells. We used the TRIzol LS Reagent (Invitrogen; REF: 10296010) to isolate the total RNAs from these groups of cells. The RNA concentrations were analyzed by the Nano-Drop OneC (Thermo Fisher Scientific). According to the manufacturer’s instructions, total RNAs were transcribed into cDNA using a high-capacity reverse transcription kit (Applied Biosystems; Lot: 2902953). The Real-time PCR reactions were performed using Power-Up SYBR Green Master Mix (Applied Biosystems; Lot: 2843446) in QuantStudio3 (Applied Biosystems). We used the online primer software Primer3 (22) to design the oligonucleotides used to amplify the host genes and other oligonucleotides used to amplify the partial BCoV-S and N genes. The relative genes and BCoV/S and BCoV/N genes expression were normalized to the β-actin according to the 2^-^^Ct^ protocol described earlier (21). The viral and host genes oligonucleotide sequences are listed in Table 2.

**Table 2:**
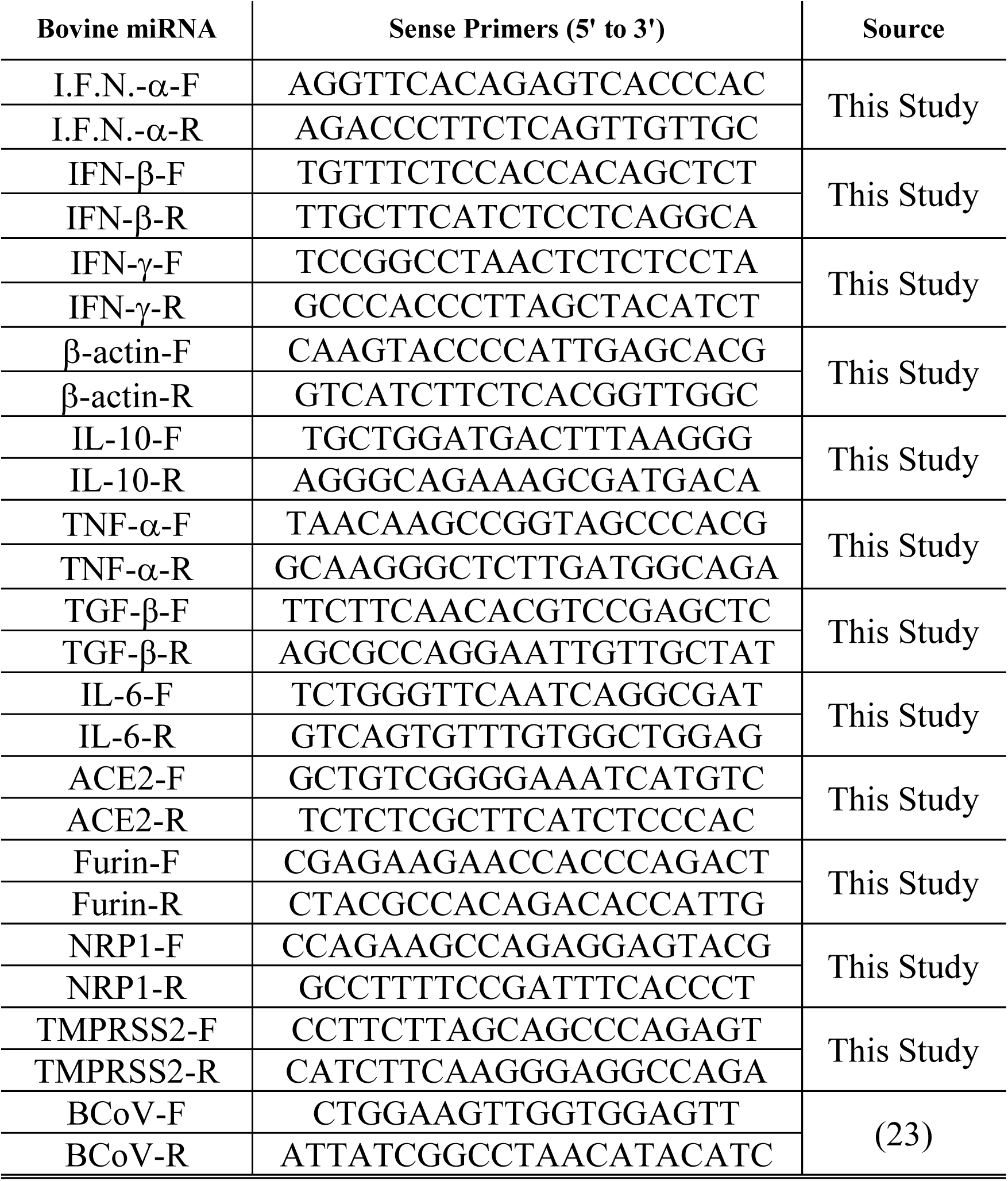
List of the oligonucleotides used for the host genes and cytokines expression profiles.

### 3.5. In Silico miRNA target gene prediction

We used the online miRNA prediction tool (TargetScan8.0) (24) to predict and identify the target genes for some candidate bovine miRNAs (bta-miRNAs). The selection criteria for the target genes include those genes showing a significant differential display among cells infected with BCoV/Ent/ BCoV/Rep or the sham-treated cells, the fold of changes of the target gene among the infected group of cells compared to the sham-infected ones, and the roles of these differentially expressed genes in the molecular pathogenesis and replication of other coronaviruses. The multiple sequence comparison of the miR-16a target site was also reported using the TargetScan prediction tools. The potential binding sites of bta-miRNA-16a targeting the BCoV genomes were predicted using three online miRNA target prediction tools, including RNA22 v2 (https://cm.jefferson.edu/rna22/), miRanda (25), and psRNA (26). The targeted site selection was based on the minimum free energy and the complementarity between the miRNA seed region and the viral genes binding sites (nucleotide 2–8 on candidate miRNAs). At least six conserved complementary nucleotides between the candidate miRNA seed region and the target location over BCoV are prerequisites to claim the potential target prediction sites.

### 3.6. Isolation of the PBMC, BCoV infection, and miRNA extraction protocols

The bovine whole blood was obtained from Lampire Biological Laboratories (LAMPIRE® Biological Labs, Inc). Briefly, the blood was aseptically withdrawn from apparently healthy animals and subjected to quality controls to ensure freedom from other pathogens and bacteria. The collected blood was mixed with 2% Ethylenediaminetetraacetic acid ferric sodium salt (Millipore-Sigma, Catalog no: E6760). The blood was processed using Histopaque-1077 (Sigma; Lot. No. RNBL7068) following the manufacturer’s instructions. The PBMCs were isolated from the buffy coat through gradient centrifugation. The isolated PBMCs were cultured in RPMI-1640 (ATCC, 30-2001) supplemented with 10% horse serum and 1% streptomycin and penicillin antibiotics and incubated at 37°C in 5% CO_2_. After 24 hours, the PBMCs were infected with (I MOI of BCoV/Ent or Resp) for two hours, followed by washing three times with sterile phosphate buffer saline (PBS) and incubating at 37°C in 5% CO_2_. The PBMCs were harvested after 72 hpi and used in the subsequent experiments. The miRNAs were extracted from the PBMCs as described above (section 3.4).

### 3.7. Transfection of miRNA Mimics (pri-miRNA) and Small interfering RNAs (siRNA)

The bta-miRNA-16a mimic sequence “UAGCAGCACGUAAAUAUUGGUG” and mirVana miRNA mimic negative control (scrambled miRNA) were purchased from Ambion, Inc. The siRNA targeting the BCoV spike glycoprotein sense sequence “CUGCUAAGAUAUAUAUGGUAUUU”, the siRNA targeting the bovine Furin sense sequence “GCACAGAGAACGACGUGGAUU”, and the si-GENOME non-targeting siRNA (scrambled siRNA) were obtained from Dharmacon™. All miRNAs and siRNAs were transfected using the Lipofectamine RNAi-MAX (Invitrogen; Ref. No. 13778-075) as described previously (27). The target cells (50 % confluence) were transfected 24hrs of subculturing with the corresponding miRNA/siRNA molecules, then infected with either BCoV/Ent, BCoV/Rep, or sham for 48 hrs after the initial transfection. As described earlier, cells that were transfected and infected were incubated for 48-72 hours post-infection (hpi). The cell culture supernatants were collected and stored at (– 80°C) for further processing, while the adherent cells were collected and processed for Western blot analysis as previously described (8, 28, 29).

### 3.8. BCoV infection protocol and the viral plaque assay

The collected cell culture supernatants from different groups of treated cells were subjected to three cycles of freezing and thawing and used for the titration of the BCoV infectivity by the plaque assay. The cell culture supernatants were incubated with the TPCK Trypsin (Thermo Fisher Scientific; REF: 20233) as described elsewhere (30, 31). An equal volume of ten ug/ml TPCK trypsin was added to each cell culture supernatants collected from variously treated cells (miRNA or siRNA treated and infected with either BCoV/Ent or BCoV/Resp independently). The cell culture supernatants containing BCoV, or sham-trypsin mixture were incubated for 30 minutes at 37°C. The MDBK cells or the human rectal tumor-18 (HRT-18) were grown in 6-well plates (1 × 10^6^) and inoculated with 10-fold serially diluted BCoV cell culture supernatants. After 1 hour incubation, aspirate supernatant and cover the cell monolayer with 3ml per well of 1.5% Sekam ME Agarose (Lonza; Cat. No. 50011), 2X EMEM (quality Biological; Cat. No. 115-073-101) supplemented with 1% Penicillin Streptomycin (Gibco; REF: 15140-122) and one ug/ml TPCK Trypsin and incubated the plate at 37°C and supplied with 5% CO_2_ for 72 three days. After 48 hrs of incubation, the cells were fixed with 4% paraformaldehyde (Thermo Fisher Scientific) overnight and stained with 1% crystal violet. The plaques were counted, and the BCoV infectivity titers per each group of treated cells were calculated using the Reed and Muench method (32).

### 3.9. Western Blot Analysis

The harvested MDBK and BEC cells from various treated groups were washed with cold PBS and then lysed with Pierce radioimmunoprecipitation assay (RIPA) lysis buffer (Thermo Fisher Scientific; REF: 89901), 1% 0.5M EDTA solution, and 1% Halt Protease & Phosphatase inhibitor (Thermo Fisher Scientific) for 5 min on ice. The collected protein samples were electrophoresed on 10% SDS-polyacrylamide gel (SDS-PAGE) and transferred to the polyvinylidene difluoride (PVDF) membrane (Bio-Rad, Cat. No. 1620177). After blocking with 5% bovine serum albumin (BSA) in Tris-buffered saline (T.B.S.) buffer containing 0.05% Tween-20 (TBST), the PVDF membranes were incubated with the respective primary antibodies followed by the incubation with the Horseradish peroxidase (HRP) conjugated secondary antibodies in the blocking solution. After three times washing with TBST, immune reactive bands were analyzed by film exposure after enhanced chemiluminescence (ECL) substrate (Bio-Rad catalog #1705060). All western blot bands were visualized by GelDoc Go Imaging System (Bio-Rad Laboratories, Inc.) and analyzed with Image J software (San Diego, US). The bovine β-actin was used to normalize the relative expression level of proteins. The primary antibodies used to detect the expression levels of the BCoV-Nucleocapsid mouse anti-bovine monoclonal (Clone: FIPV3-70; Cat. No. MA1-82189), BCoV-Spike rabbit anti-bovine polyclonal (Cat. No. PA5-117562), and β-actin rabbit anti-bovine polyclonal (Cat. No. PA1-46296) were purchased from Invitrogen. The Furin rabbit anti-bovine polyclonal (Cat. No. ARP45328_P050), the ACE2 rabbit anti-bovine polyclonal (Cat. No. ARP53751_P050), the TMPRSS2 rabbit anti-bovine polyclonal (Cat. No. ARP46628_P050), and the anti-NRP1 rabbit anti-bovine polyclonal (Cat. No. ARP59101_P050) were purchased from Aviva Systems Biology. The corresponding secondary antibodies for each protein were used, including the horseradish peroxidase (H.R.P.)-conjugated IgG (H+L), goat anti-rabbit (REF: 31460), and goat anti-mouse (REF: 31430) were obtained from Invitrogen. The western blot band density was measured using ImageJ software (33). The mean band density of each protein was normalized by dividing its values by the mean band density of the housekeeping gene (B-actin) for each sample separately (33).

### 3.10. Cloning of the 3’UTR of the bovine Furin and the mutated seed region of bta-miRNA-16a in the dual luciferase expression vector and the Dual Luciferase Assay

The Wild-Type (WT) and Mutant (Mut) binding sites of bta-miRNA-16a within the 3’UTR of the bovine Furin were synthesized by a commercial provider (GenScript USA Inc). Briefly, the WT Furin-3’UTR construct region spanning the binding region of bta-miRNA-16a was annealed and inserted between the *SacI* and *XhoI* region of pmirGLO Dual-Luciferase miRNA Target Expression Vector (Promega, Catalog number: E1330). Mutations within the bta-miRNA16a WT construct were generated by PCR-based site-directed mutagenesis. The binding region of Furin with bta-miRNA-16a was mutated from “5’-TGCTGCT-3’” to “5’-GATGATC-3’” according to the manufacturer’s instructions. Both constructs were confirmed by the restriction enzyme digestion and sequencing (S1 Fig). Per the manufacturer’s instructions, the dual luciferase assay was conducted using the Pierce Renilla-Firefly Luciferase Dual Assay Kit (Thermo Scientific; Ref. No. 16185). To determine the expression levels of Furin in the bta-miRNA-16a transfected MDBK and HEK 293T cells, we plated the cells into 24-well plates and co/transfect these cells with 100 ng of bta-miRNA-16a mimic and 100 ng of either pmiR-GLO-Furin-WT or pmiR-GLO-Furin-Mut plasmids using Lipofectamine 3000 (Invitrogen Catalog # L3000001) transfection reagent. An empty plasmid (pmiR-GLO) transfected with the bta-miRNA-16a, and pmiR-GLO-Furin-WT transfected with scrambled miRNA were used as negative controls. All experiments were repeated three independent times.

### 3.11. Statistical analysis

All the reported results in this study were displayed as means ± S.D., analyzed with GraphPad Prism v9. One-way analysis of variance (ANOVA) with Tukey’s or Dunnett’s test was performed among multiple groups. The Student’s t-test was used for the paired comparison among the samples. The P values < 0.05 were considered statistically significant. Statistical significance in the figures is indicated as follows: * p < 0.05, ** p < 0.01, *** p < 0.001, *** p < 0.0001, and ns, not significant. Data were combined from at least three independent experiments unless otherwise stated.

## 4. Results

### 4.1. Bovine Coronavirus (BCoV) Infection Induces Differential Display of the Host Cell mRNA and miRNA Expression Profiles

The Madine Darby Bovine Kidney (MDBK) and Bovine Vascular Endothelial cells (BEC) were infected with Bovine coronavirus Enteric (BCoV/Ent) and Bovine coronavirus Resp (BCoV/Resp) isolates at a multiplicity of infection (MOI) of 1. The cytopathic effect (C.P.E.) and morphological changes were observed in the infected cells at 72 hours post-infection (hpi) (Fig 1A, 1B). The morphological observations of the BCoV-infected cells revealed a more prominent CPE in the BCoV/Ent isolate-infected groups compared to the BCoV/Resp isolate-infected groups, especially in MDBK cells (Fig 1A). The q-RT-PCR results demonstrated that BCoV/Ent and BCoV/Resp isolates successfully infected and propagated on both MDBK and BEC cells (Fig 1C, 1D). The magnitude of the BCoV genomic viral load was more prominent in the BCoV/Ent infected groups compared to the BCoV/Resp infected cells (Fig 1C, 1D). The KEGG pathway enrichment analysis analyzed from NGS data revealed significant virus-related pathways involved in response to BCoV/Ent and BCoV/Resp infection (Fig 1E, 1F). The KEGG pathway enrichment analysis of the BCoV/Ent-infected group showed significant gene count numbers in human cytomegalovirus (hCMV) infection and Hepatitis B (Fig 1E). The KEGG pathway enrichment analysis in the BCoV/Resp infected group reflects the highest significance to the coronavirus COVID-19 pathway, followed by hCMV and hepatitis B (Fig 1F). The NGS analysis revealed more differentially expressed host genes in the BCoV/Resp group (Fig. 1E). At the same time, miRNA expression patterns were consistent across both BCoV/Ent and BCoV/Resp groups (Fig 1F). These results highlight distinct infection patterns between BCoV/Ent and BCoV/Resp isolates in MDBK and BEC cells. Differential expression of host genes involved in various viral infections indicates a broad immune response.

**Fig 1:**
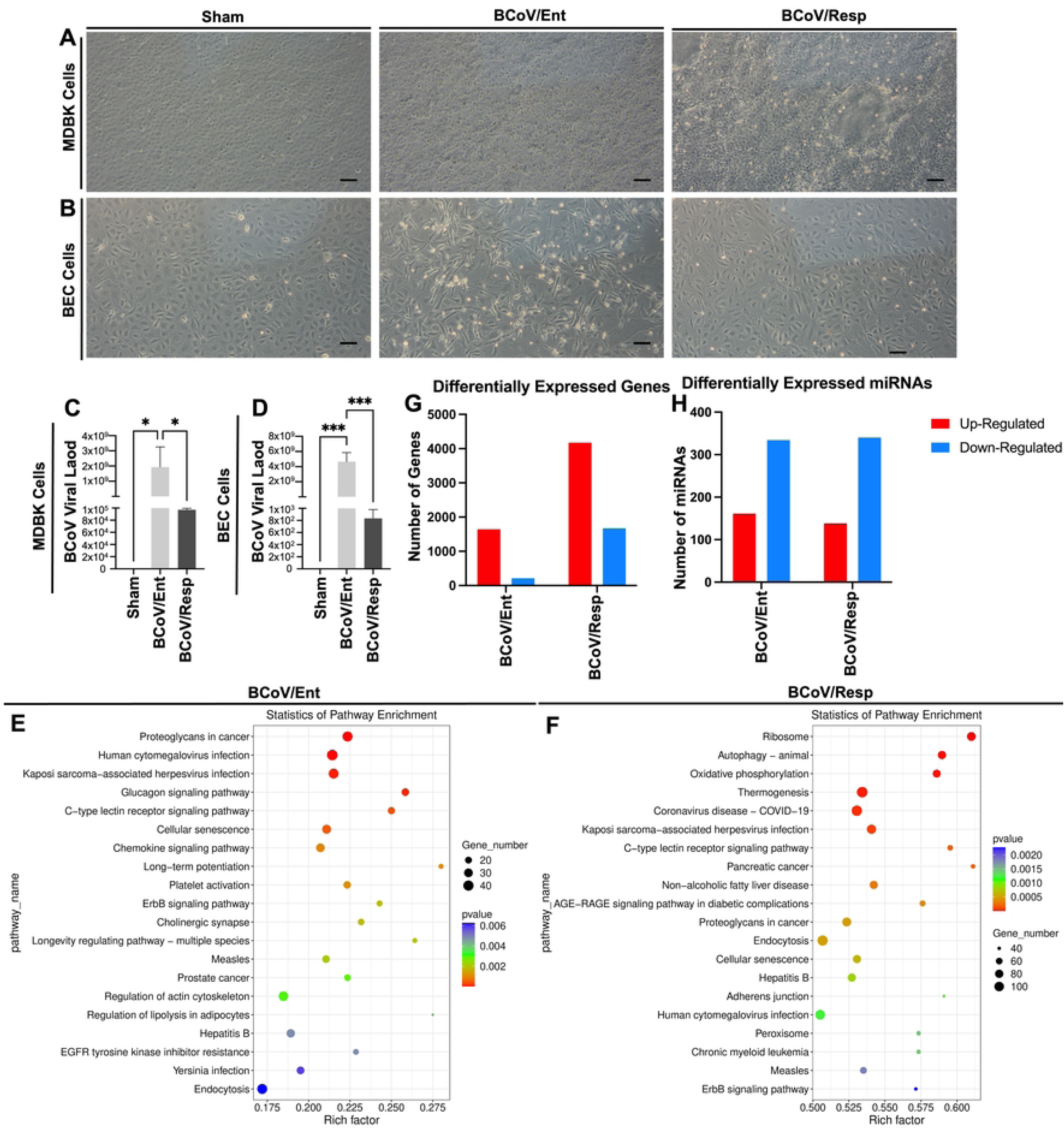
The host cell miRNAs expression profiles during Bovine Coronavirus (BCoV) replication. **(A)** Morphological observation of MDBK cells in control (Sham), cytopathic effect (C.P.E.) at 72 hours post-infection (hpi) with BCoV enteric (BCoV/Ent) and respiratory (BCoV/Resp) isolate (10x magnification). **(B)** Morphological observation of BEC cells: Cl, C.P.E. at 72 hpi with BCoV/Ent and BCoV/Resp isolates (10x magnification). **(C)** The qRT-PCR analysis of BCoV genomic viral load in MDBK cells. **(D)** The qRT-PCR analysis of BCoV genomic viral load in BEC cells. **(E)** KEGG pathway analysis from Next-Generation Sequencing (NGS) data indicates differentially expressed host cellular pathways during infection with the BCoV/Ent isolate and **(F)** during BCoV/Resp isolate. **(G)** NGS data indicating the number of differentially expressed genes in BCoV/Ent and BCoV/Resp infected groups. **(H)** NGS data illustrating the number of differentially expressed miRNAs in BCoV/Ent and BCoV/Resp infected groups.

### 4.2. The bta-miRNA-16a is differentially expressed in target cells during the BCoV Ent/Resp infection

We analyzed the data obtained from the NGS experiments of the BCoV-infected cells to identify the potential miRNAs showing a differential expression in the BCoV/Ent and BCoV/Resp infected groups (Fig 2A). Our results showed that the bta-miRNA-16a was down-regulated in the BCoV/Ent group and upregulated in the BCoV/Resp groups compared to the sham-infected cells (Fig 2A). In the case of the BCoV-infected MDBK cells, the bta-miRNA-16a was significantly upregulated in the BCoV/Resp group, while no significant expression of bta-miRNA-16a was observed in the BCoV/Ent group (Fig 2B). In the case of the BCoV-infected BEC cells, bta-miRNA-16a was down-regulated up to 1.3-fold in the BCoV/Ent group and upregulated up to 1.74-fold in the BCoV/Resp group (Fig 2C). The bovine PBMCs showed significant down-regulation of bta-miRNA-16a up (2.8-folds) in the BCoV/Ent infected group, but no substantial changes in the beta-miRNA-16a expression were observed in the BCoV/Resp group (Fig 2D). These results demonstrate consistent expression patterns between NGS and qRT-PCR analyses, validating the differential expression pattern of bta-miRNA-16a in BCoV/Ent and BCoV/Resp infected groups.

**Fig 2:**
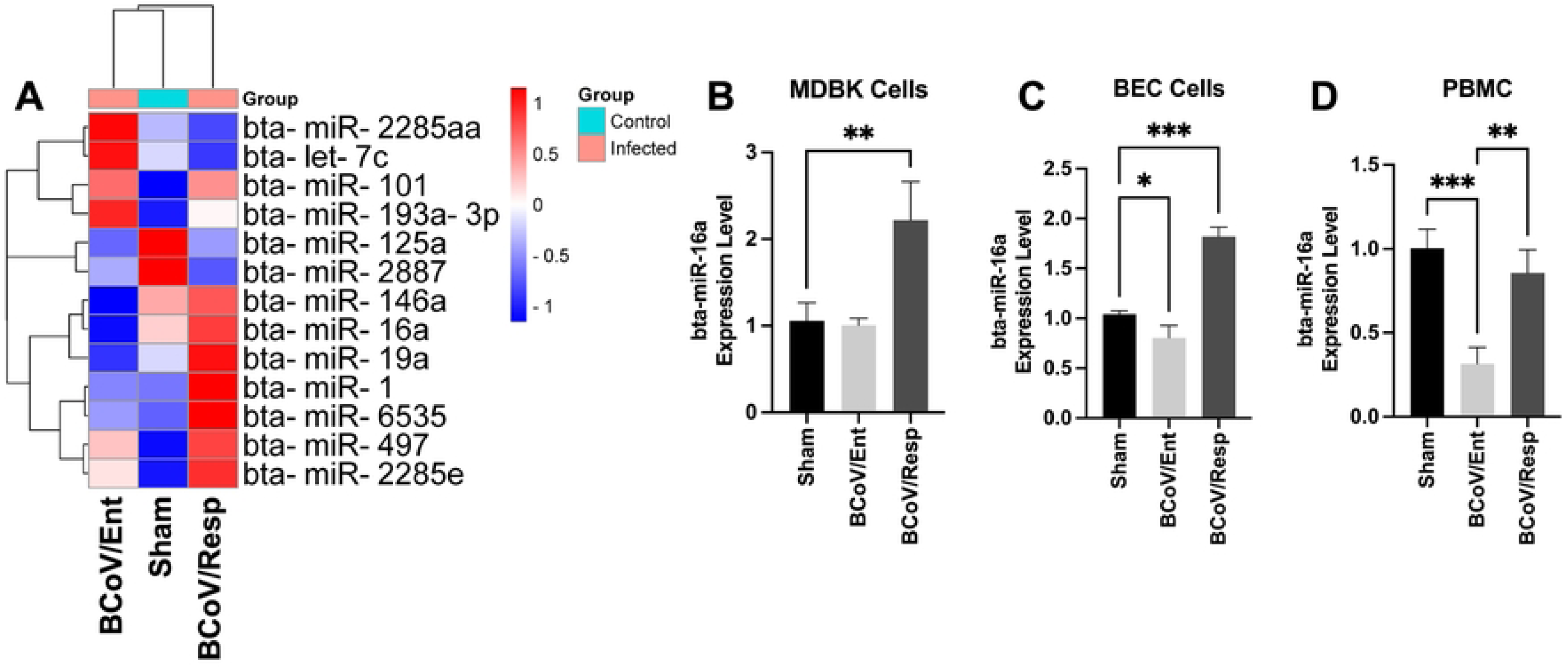
Bovine miRNA expression profile during BCoV/Ent or BCoV/Resp infection in bovine cell lines. **(A)** Heatmap showing the differentially expressed miRNAs in the sham, BCoV/Ent, and BCoV/Resp infected group of cells based on the NGS data analysis. **(B)** The qRT-PCR data shows the expression levels of bovine miR-16a in MDBK cells. **(C**) The qRT-PCR data shows the expression levels of bovine miR-16a in BEC cells. **(D**) The qRT-PCR data shows the expression levels of bovine miR-16a in PBMC isolated from bovine blood. The heat map was plotted using (https://www.bioinformatics.com.cn/en), a free online data analysis and visualization platform.

### 4.3. The bta-miRNA-16a restricts BCoV replication

The in-silico analysis revealed that bta-miRNA-16a has multiple targeting sites across the BCoV genome, including one target in the ORF1b and two other targets within the BCoV-spike glycoprotein (Table 3). To further validate this prediction, the bta-miRNA-16a was transfected into MDBK and BEC cells independently, followed by BCoV/Ent/Resp infection independently in parallel to the scrambled miRNA (miRNA-Scr). The qRT-PCR analysis revealed significant up-regulation of bta-miRNA-16a in the miRNA-16a transfected group compared to miRNA-Scr in the MDBK and the BEC cells (Fig 3B, 3C). Our data shows that bta-miRNA-16a expression was down-regulated after BCoV infection in non-transfected cells (Fig 3B, 3C). Our results also showed marked inhibition on the BCoV genome expression level in bta-miRNA-16a transfected groups compared to the miRNA-Scr transfected MDBK and BEC cells (Fig 3D, 3E). The viral plaque assay revealed up to 1.24-fold inhibition in the BCoV infectivity in the bta-miRNA-16a transfected group of cells (Fig 3F). On the protein levels, the western blot analysis showed significant inhibition of BCoV-Nucleocapsid (BCoV-N) and BCoV-Spike (BCoV-S) proteins after bta-miRNA-16a transfection and BCoV infection in MDBK cells (Fig 3G, 3H, and 3I). The BEC cells showed marked inhibition of the BCoV-N and BCoV-S protein expression levels (Fig 3J, 3K, and 3L). The findings suggest that bta-miRNA-16a targets the spike protein of BCoV, playing a crucial role in restricting BCoV replication.

**Fig 3:**
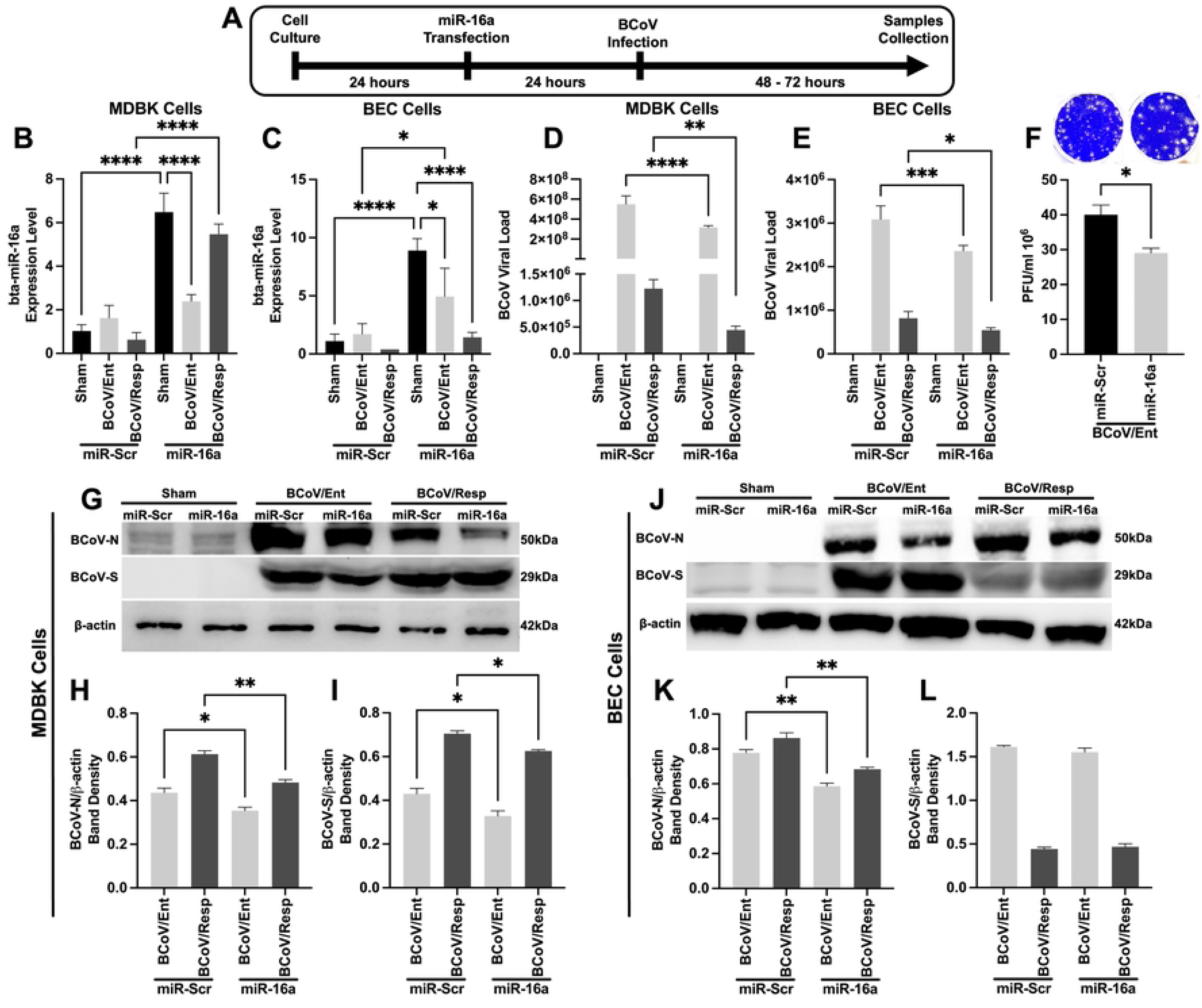
Bovine miR-16a overexpression markedly inhibits BCoV replication and virus infectivity. **(A)** A Schematic representation of miR-16a transfection and BCoV infection experiments; Transfection of miR-Sc and miR-16a into MDBK and BEC cells, followed by infection with BCoV enteric and respiratory isolates for 48 to 72 hours, and subsequent sample collection for further analysis. **(B)** miR-16a expression in MDBK cells: qRT-PCR data displaying the expression pattern of bovine miR-16a in scrambled and miR-16a transfected MDBK cells. **(C)** miR-16a expression in BEC. cells: qRT-PCR analysis showing the expression pattern of bovine miR-16a in scrambled and miR-16a transfected BEC cells. **(D)** BCoV genomic level in MDBK cells: qRT-PCR analysis demonstrating genome level of BCoV in scrambled and miR-16a transfected MDBK cells. **(E)** BCoV genomic level in the BEC cells: qRT-PCR analysis illustrating genome level of BCoV in scrambled and miR-16a transfected BEC cells. **(F)** BCoV enteric infectivity level in MDBK cells: Viral plaque assay indicating the infectivity level of BCoV enteric isolate in scrambled and miR-16a transfected the MDBK cells. **(G)** Protein expression in MDBK cells: Western blot analysis of BCoV-nucleocapsid (BCoV-N) and BCoV-spike (BCoV-S) observed in MDBK cells transfected with scrambled and miR-16a. **(H)** Western blot band density of BCoV-N protein normalized with β-actin in MDBK cells. **(I)** Western blot band density of BCoV-S protein normalized with β-actin in the MDBK cells. **(J)** BCoV protein expression in BEC cells: The BEC. cells were transfected with miRNA-Sc and bta-miRNA-16a, and the western blot analysis assessed protein expression of the BCoV-N and the BCoV-S protein. **(K)** Western blot band density of BCoV-N protein normalized with β-actin in BEC cells. **(L)** Western blot band density of BCoV-S protein normalized with β-actin in BEC cells.

**Table 3:**
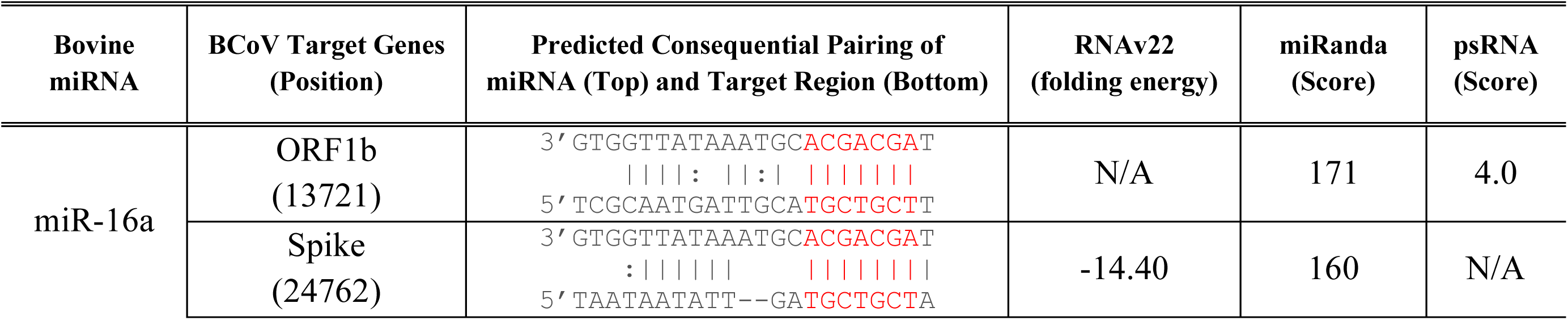
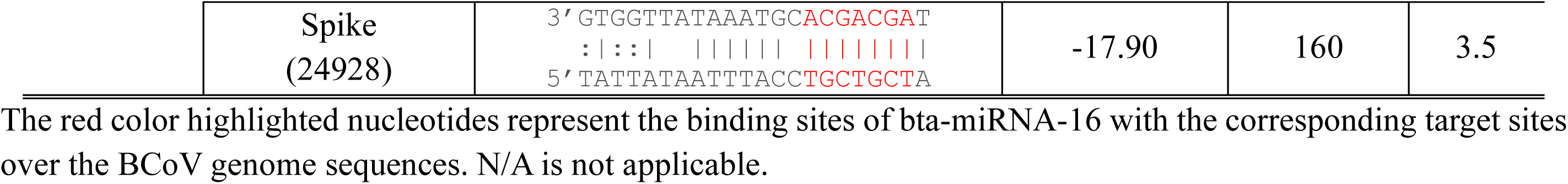
Online prediction of the host bta-miRNA-16a targeting BCoV genome at multiple locations.

### 4.4. The bovine Furin is a novel target for the host bta-miRNA-16a

To further evaluate the mechanism of action of bta-miRNA-16a-based inhibition of BCoV replication, the in-silico prediction was conducted between the bta-miRNA-16a, and some potential target host genes involved in BCoV infection. The binding region of Furin is conserved across different mammalian species, including humans and mice (Fig 4A). The online bioinformatical tool “TargetScan” identified that bta-miRNA-16a could potentially target the 3’UTR of the bovine host cellular Furin (Fig 4B). Therefore, to examine the effect of bta-miRNA-16a on the expression levels of the host Furin on the mRNA and the protein levels, the bta-miRNA-16a was transfected into both MDBK and BEC cells, and the inhibition of host cellular Furin was observed. The mRNA of host cellular Furin was down-regulated up to 70 percent in the bta-miRNA-16a transfected groups compared to the miRNA-Scr in MDBK cells (Fig 4C). The protein level of Furin was down-regulated by approximately 30 percent in the bta-miRNA-16a transfected groups compared to the miR-Scr in MDBK cells (Fig 4D, 4E). In the BEC cells, up to 50 percent down-regulation of host furin protein was observed in miRNA-16a transfected groups (Fig 4G, 4H). However, no significant down-regulation of furin mRNA in the BEC cells transfected group of the miRNA-Scr transfected cells (Fig 4F). To validate Furin as a potential novel target of bta-miRNA-16a, a dual-luciferase reporter plasmid containing the 3’UTR region of Furin containing the binding region of bta-miRNA-16a was constructed (Fig 4I). The mutant Furin was generated by site-directed mutagenesis of the bta-miRNA-16a binding region (Fig 4I). The successful insertion of 3’UTR of Furin and the mutant construct was validated by the double digestion with *XhoI* and *SacI* restriction enzymes, followed by gel-based PCR and nucleotide sequencing (Fig S1A – S1D). The wild-type and mutant pmiR-GLO luciferase construct was co-transfected with miR-16a or scrambled miRNA into HEK cells. The dual-luciferase assay revealed that co-transfection of wild-type Furin with miRNA-16a showed a significant reduction of more than 50 percent of the relative luciferase activity compared to the miRNA-Scr co-transfected with wild-type Furin or the bta-miRNA-16a co-transfected with mutated Furin (Fig 4J). The overall results confirm that the bta-miRNA-16a can effectively target and significantly down-regulate host cellular Furin.

**Fig 4:**
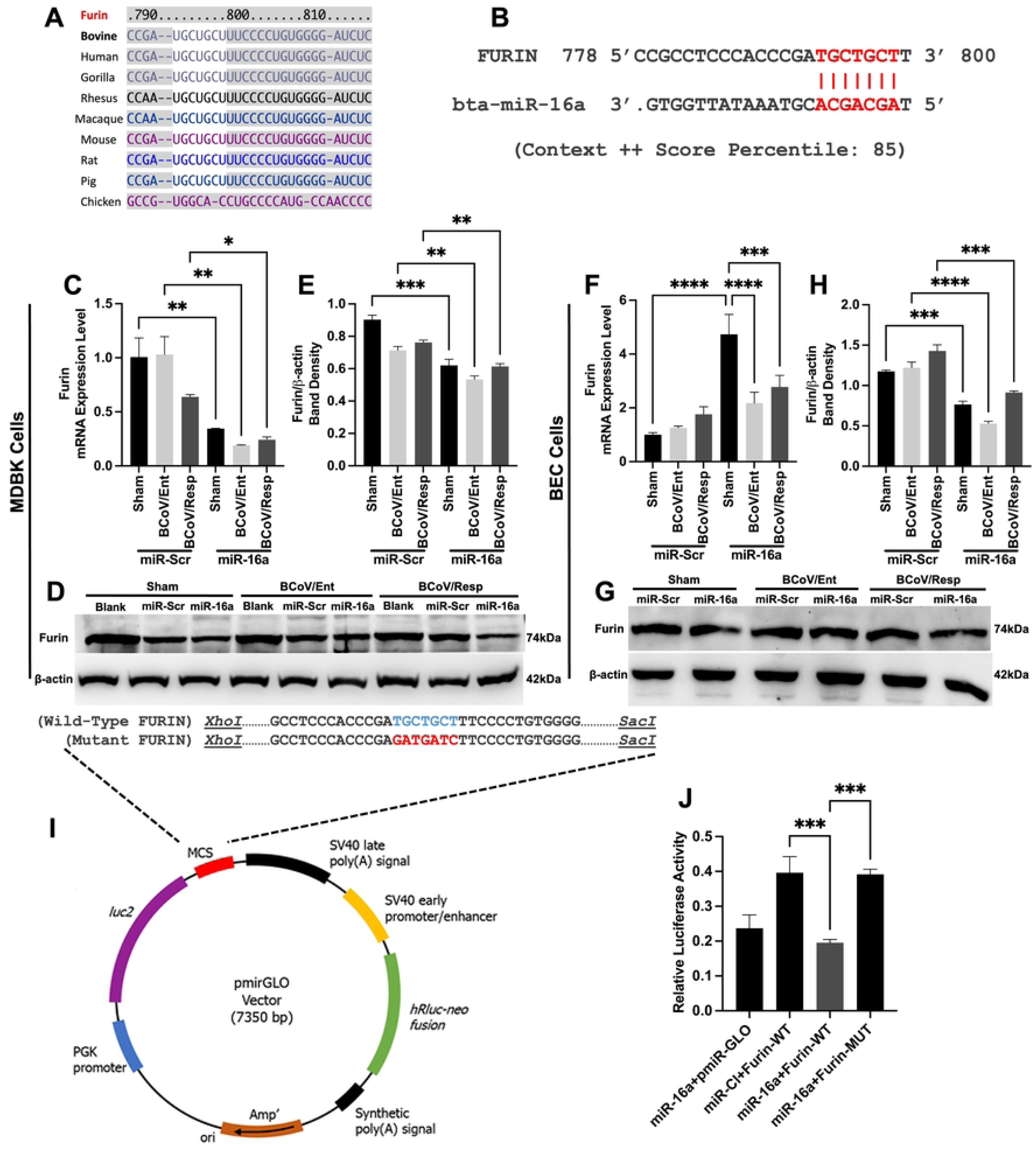
Host cell Furin as a novel target for bta-miRNA-16a and the impact of this targeting on BCoV replication. **(A)** The multiple sequence alignment shows that the bta-miRNA-16a/Furin binding region is conserved among eight species, including humans, mice, and bovine. **(B)** In-silico prediction of the bta-miRNA-16a targeting the 3’ UTR of the host cell Furin. The context ++ score percentile represents the binding energy of miRNA with the target gene. **(C)** The MDBK cells were transfected with either the miR-Scr or miR-16a, followed by BCoV/Ent or BCoV/Resp isolates infection. After 72 hours, samples were tested by the qRT-PCR. Data analysis assessed the mRNA expression level of host Furin in both Sc-miRNA or miR-16a transfected groups of cells. **(D)** Western blot analysis of host Furin protein expression in miR-Scr and miR-16a transfected groups in MDBK cells. **(E)** Western blot band density of Furin protein normalized with β-actin in MDBK cells. **(F)** The BEC cells were transfected with scrambled and miR-16a, followed by BCoV enteric and respiratory isolates infection. After 72 hours, samples were collected for qRT-PCR analysis to assess the mRNA expression level of host Furin in both scrambled and miR-16a transfected groups. **(G)** Western blot analysis of host Furin protein expression in scrambled and miR-16a transfected groups in BEC cells. **(H)** Western blot band density of Furin protein normalized with β-actin in BEC cells. **(I)** Diagram of luciferase reporter plasmids. A fragment of the 3’UTR of Furin wild-type (W.T.) carrying miR-16a binding site (indicated in blue) was inserted into the *XhoI* and *SacI* restriction enzymes sites of the pmiR-GLO luciferase vector. The seed region of Furin was mutated by site-directed mutagenesis to produce mutant (M.U.T.) Furin (indicated in red). **(J)** The results of the dual-luciferase reporter assay. H.E.K. cells were transfected with different combinations of the Furin 3’UTR (W.T.) or mutant (Mut) plasmid or vector only (pmir-GLo) with miR-16a and miR-Scr, as indicated. The dual Luciferase assay was conducted, and the relative luciferase activity was calculated using red firefly luciferase signals as a normalization control for the green Renilla luciferase signals.

### 4. 5. Applying siRNA targeting the BCoV-Spike Glycoprotein and the Host Furin Inhibits BCoV Replication on the gene and protein expression levels

To confirm the action of bta-miRNA-16a targeting the BCoV-S proteins and the cellular host cell Furin on BCoV replication., we designed two sets of siRNAs. The first siRNA targets the bta-miRNA-16a target site within the BCoV/S glycoprotein. The second siRNA targets the bta-miRNA-16a target in the 3’UTR of the host cell Furin. Bovine cells were transfected with siRNAs, followed by independent infection of BCoV/Ent or BCoV/Resp isolates. Samples were collected after 72 hpi for subsequent analysis (Fig 5A). The MDBK cells were transfected with different concentrations of siRNAs (50ng and 100ng). The results revealed a gradual reduction in bovine host Furin protein levels with increasing siRNA-Furin concentration (Fig 5B, 5C). Similarly, the BCoV-spike protein exhibited a similar inhibition in the expression levels with increasing the siRNA-spike concentration (Fig 5D, 5E), confirming the efficacy of siRNAs in suppressing the production of BCoV-S and host Furin. The mRNA expression level of host cellular Furin was also significantly down-regulated in the Furin-siRNA treated groups compared to Sc-siRNA in both the MDBK and the BEC. cells (Fig 5F, 5G). Following the silencing of Furin and spike, we examined their impact on BCoV replication. Results revealed approximately 58 percent (≈2.39-fold) inhibition in the BCoV genomic level in siRNA-Furin transfected MDBK cell groups (Fig 5H). In BEC cells, the inhibitory actions of the applied siRNAs were even more pronounced, with about 75 percent (≈4.26-fold) inhibition in the genomic level of BCoV (Fig 5I). The inhibition of BCoV replication was particularly prominent in BCoV/Ent-infected groups. The silencing of the BCoV-S glycoprotein demonstrated about 61 percent (≈2.61 fold) inhibition in BCoV/Ent and about 74 percent (≈3.85 fold) inhibition in BCoV/Resp groups in the BEC-treated cells (Fig 5K).

**Fig 5:**
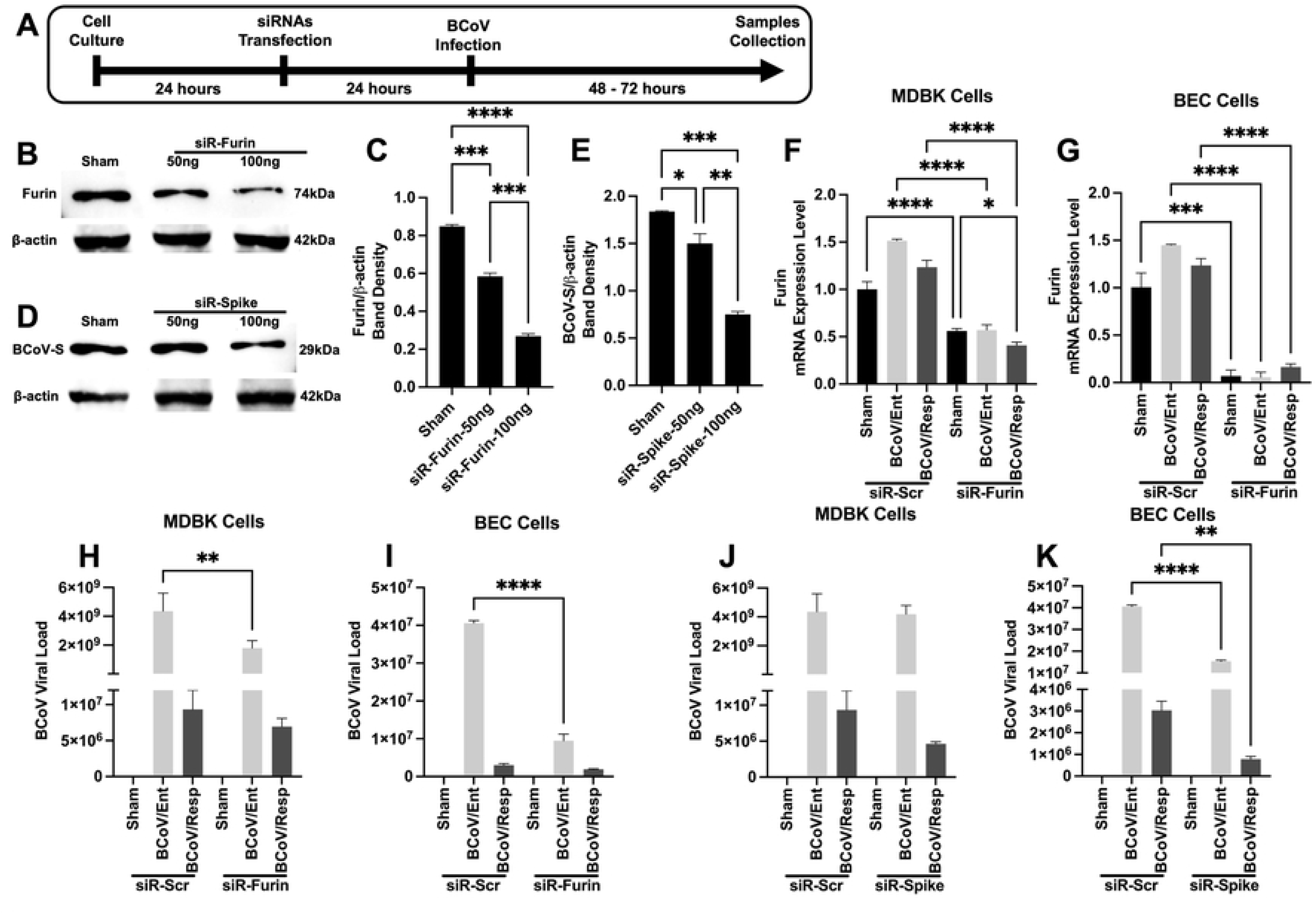
The impacts of silencing of host Furin and BCoV-S glycoprotein on the BCoV replication. **(A)** Scheme of siRNA transfection experiment: Illustrating the process of transfecting scrambled, siRNA-Furin, and siRNA-BCoV-S glycoprotein into the MDBK and the BEC cells, followed by infection with BCoV/Ent or BCoV/Resp isolates for 48 to 72 hours, and subsequent sample collection for further analysis. **(B)** The MDBK cells were transfected with siRNA-Furin at 50ng and 100ng, followed by western blot analysis for Furin protein expression. **(C)** Western blot band density of Furin protein normalized to the β-actin in the MDBK cells. **(D)** The MDBK cells were transfected with siRNA-S at 50ng and 100ng, followed by western blot analysis for BCoV spike protein expression. **(E)** Western blot band density of BCoV spike protein normalized with β-actin in MDBK cells. **(F)** qRT-PCR analysis showing Furin expression in scrambled and siRNA-Furin transfected MDBK cells **(G)** and BEC cells. **(H)** The qRT-PCR analysis demonstrating the genomic level of BCoV in scrambled and siRNA-Furin transfected the MDBK cells **(I)** and the BEC cells. **(J)** qRT-PCR analysis illustrating genome level of the BCoV in scrambled and siRNA-BCoV-Spike transfected MDBK cells; **(K)** and BEC cells.

### 4. 6. Applying siRNA of the BCoV-S glycoprotein and the host cell-furin Inhibits the BCoV infectivity in Host Cells

The viral plaque assay was conducted to evaluate the impacts of siRNA-Furin and siRNA-BCoV-spike glycoprotein on the infectivity of the BCoV particles. Following transfection with siRNA-Furin and siRNA-BCoV-spike, the MDBK cells were infected with BCoV, and cell lysates were analyzed to assess BCoV inhibition at the post-transcriptional level. Results showed about 1.9-fold inhibition in the infectivity level of BCoV enteric isolate after silencing host Furin (Fig 6A). Silencing BCoV Spike in MDBK cells resulted in about 1.47-fold inhibition in BCoV infectivity level (Fig 6B). In the MDBK-treated cells, the host Furin protein expression level was inhibited by about 50 percent after silencing host Furin and by about 42 percent after silencing the BCoV spike glycoprotein (Fig 6C, 6D). Results revealed about 2.5-fold inhibition of BCoV spike protein in siRNA-Furin and about 1.38-fold inhibition in siRNA-spike transfected group compared to the scrambled (Fig 6C, 6E). In the host Furin silenced group of cells, the BCoV nucleocapsid protein was inhibited about 1.79-fold and 1.97-fold in BCoV/Ent and BCoV/Resp infected groups, respectively (Fig 6C, 6F). Silencing of the BCoV spike inhibited BCoV nucleocapsid protein about 1.24-fold and 1.31-fold in BCoV/Ent and BCoV/Resp infected groups, respectively (Fig 6C, 6F). Notably, the downregulation of spike, nucleocapsid, and Furin was consistent in control and BCoV-infected groups, indicating that siRNA-spike and siRNA-Furin inhibit protein production at the post-transcriptional level, independent of BCoV infection.

**Fig 6:**
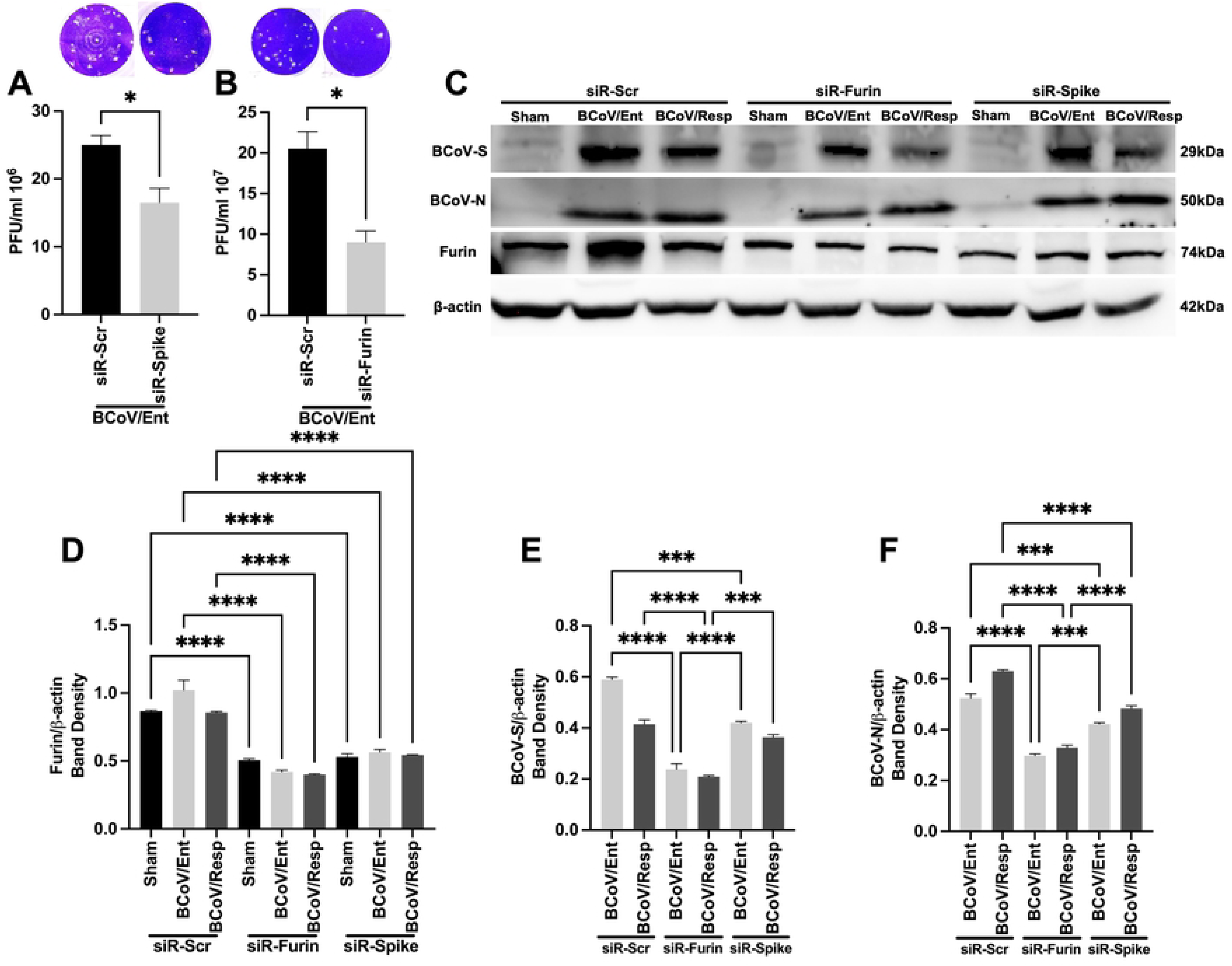
Silencing of the host cell Furin and BCoV-S glycoprotein negatively impacted the BCoV infectivity in cells. **(A)** The BCoV/Ent or BCoV/Resp viral particle infectivity is inhibited in cells transfected with siRNA-Spike, or (B) siRNA-Furin transfected groups as measured by the viral plaque assay. The plaque-forming unit (PFU) counted the virus titer in each virus-infected group. **(C)** The MDBK cells were transfected with scrambled siRNA (Scr-siRNA), siRNA-Furin, and siRNA-Spike, followed by infection of BCoV/Ent and BCoV/Resp isolate. Cells were lysed after 72 hpi, and western blot analysis was performed to observe the protein expression of BCoV-S, BCoV-N, and host Furin proteins. **(D)** Western blot band density of Furin protein; **(E)** BCoV-spike; **(F)** and BCoV-N protein was normalized to the β-actin in MDBK cells.

### 4. 7. The bta-miRA-16a modulates the expression levels of some essential Coronavirus replication-related proteins (ACE2, NPR1, TMPRSS2) during BCoV replication

Herein, we evaluate the possible effect of bta-miRNA-16a on other host cell surface receptors involved in some other coronavirus infections, particularly the severe acute respiratory syndrome coronavirus (SARS-CoV-2). The MDBK cells were transfected with miRNA-Scr and the bta-miRNA-16a and monitored for the mRNA and the protein expression levels of some selected host cell proteins (Angiotensin-converting enzyme 2 (ACE2), transmembrane protease, serine 2 (TMPRSS2), and Neuropilin-1 (NRP1)). Results revealed that the ACE2 protein expression level was inhibited by about 22 percent in the bta-miRNA-16a transfected groups compared to the miRNA-Scr transfected cells (Fig 7A, 7C). In the bta-miRNA-16a transfected group, the mRNA of ACE2 was downregulated up to 50 percent after BCoV infection (Fig 7B). The host NRP1, mRNA, and protein expression markedly upregulated after BCoV infection in the miRNA-Scr transfected cells, but it showed down-regulation in the bta-miRNA-16a transfected group of cells, particularly after BCoV infection (Fig 7A, 7D, 7E). The mRNA of TMPRSS2 also showed down-regulation after BCoV infection in the bta-miRNA-16a transfected group of cells (Fig 7F); however, no significant changes in protein expression levels between miRNA-Scr and the bta-miRNA-16a transfected groups was observed (Fig 7A, 7G).

**Fig 7:**
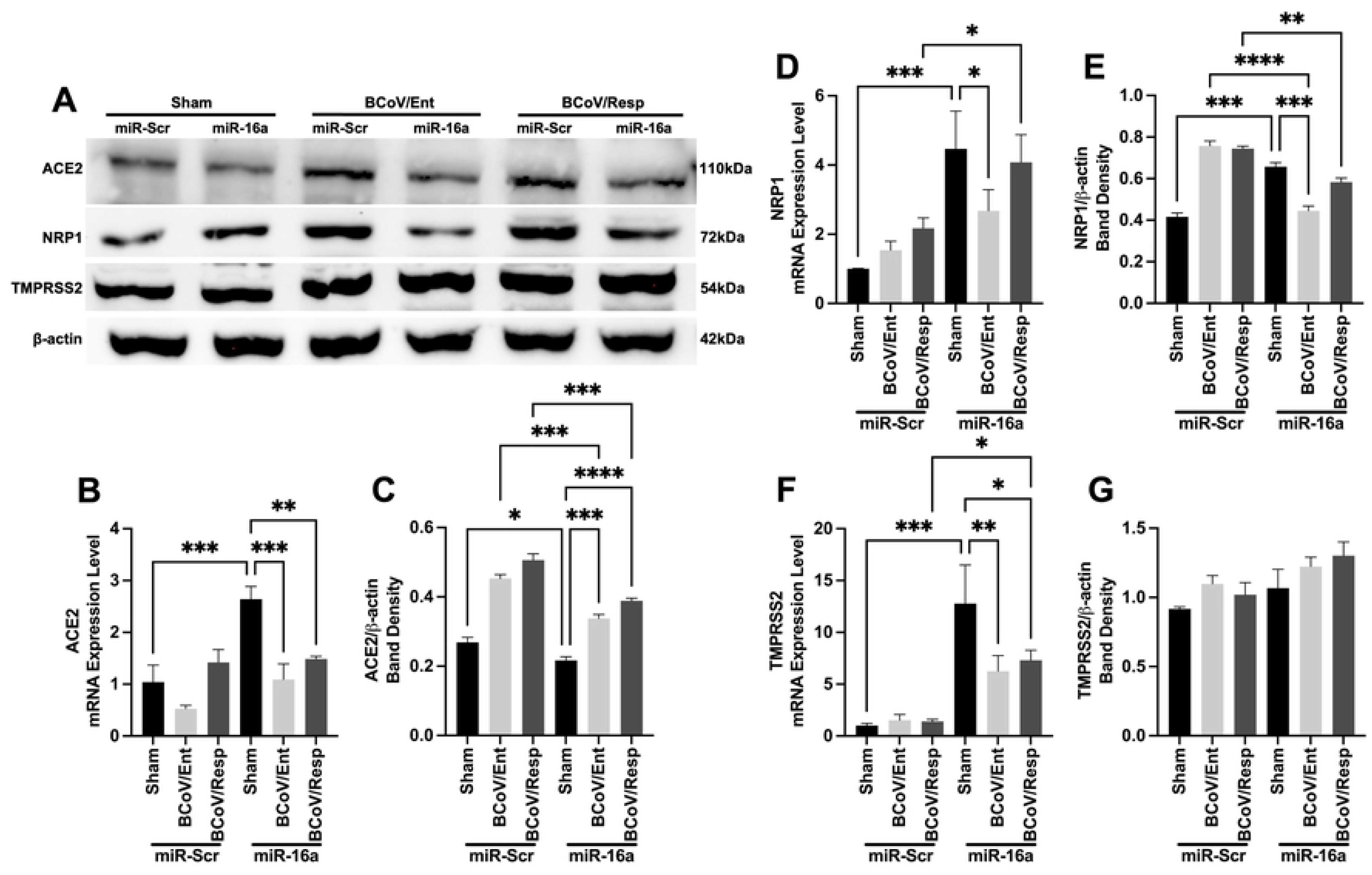
The impact of host miR-16a overexpression on some selected SARS-CoV-2 replication-related proteins (ACE2, NRP1, and TMPRSS2). **(A)** The MDBK cells were transfected with either the Scr-miRNA or the bta-miR-16a, followed by BCoV/Ent or BCoV/Resp isolates infection. Both the cell culture supernatants and the cell lysate were collected after 72-hpi for the qRT-PCR and western blot analysis of host ACE2, NRP1, TMPRSS2, and β-actin. **(B)** Results of the qRT-PCR analysis of host ACE2 mRNA expression in the Scr-miRNA and bta-miRNA-16a transfected groups. **(C)** Results of the Western blot band density of ACE2 protein normalized to the β-actin in MDBK cells. **(D)** Results of the qRT-PCR analysis of the host NRP1 mRNA expression levels in various groups of cells. **(E)** Results of the Western blot band density of NRP1 protein normalized to the β-actin in MDBK cells. **(F)** Results of the qRT-PCR analysis of host the host TMPRSS2 mRNA expression levels. **(E)** Results of the Western blot band density of TMPRSS2 protein normalized with β-actin in MDBK cells.

### 4. 8. Inhibition of Host Furin expression through bta-miRNA-16 targeting potentially activates other alternative Pathways to rescue the BCoV Infection in Host Cells

To further explore the roles of the host cell Furin in the BCoV infection, two siRNAs were used to inhibit BCoV regulation by targeting the BCoV-S glycoproteins and the bta-miRNA-16 target sites within the 3’UTR of the host cell Furin. We independently applied those two siRNAs through transfection into bovine cells and investigated the mechanism of roles of the bovine ACE2, TMPRSS2, and NRP1 during BCoV/Ent/Resp isolates infection. The siRNA-BCoV-S and siRNA-Furin were transfected into the MDBK cells, and the mRNA and protein expression of the host ACE2, TMPRSS2, and NRP1 were assessed using qRT-PCR and Western blot using the relevant antibodies, respectively. The TMPRSS2 expression did not show any significant alteration in their expression in the case of the siRNA-Furin or siRNA-BCoV-S transfected group of cells (Fig 8A, 8B, 8C). The host ACE2 mRNA and protein expression levels were significantly down-regulated after siRNA-Furin transfection (Fig 8A, 8D, 8E). The ACE2 protein expression was also down-regulated in the siRNA-spike transfected group but only after BCoV infection (Fig 8A, 8D, 8E). The NRP1 mRNA and protein expression showed a similar down-regulation pattern in siRNA-Furin transfected groups, while it was only down-regulated after BCoV infection in the siRNA-spike transfected group (Fig 8A, 8F, 8G). The mRNA expression of both ACE2 and NRP1 down-regulated after BCoV infection in the siRNA-BCoV-S transfected group, suggesting their involvement in BCoV infection. These findings mirror the outcomes of bta-miRNA-16a transfection, suggesting a direct correlation between ACE2 and NRP1 expression with host Furin expression. In contrast, TMPRSS2 appears to operate independently of host Furin upon BCoV infection.

**Fig 8:**
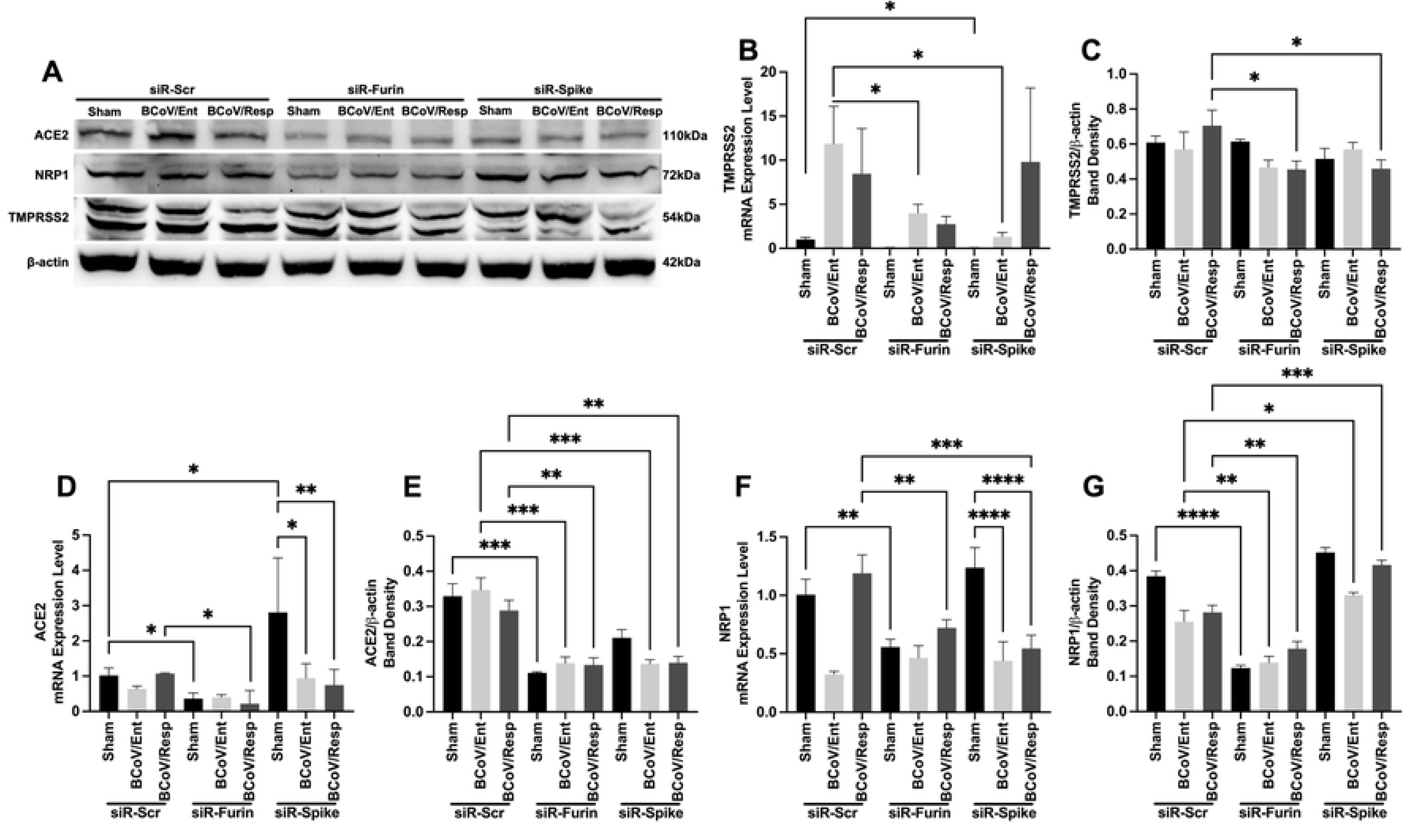
Silencing of the host cell Furin and BCoV-S glycoprotein impacted the expression levels of some SARS-CoV replication-related proteins on the mRNA and protein levels, which further impacted BCoV/Ent/BCoV/Resp replication. **(A)** The MDBK cells were transfected with either Scr-siRNA control, siRNA-Furin, or siRNA-Spike, followed by either BCoV/Ent or BCoV/Resp isolates infection. Cell lysates were collected after 72 hpi and subjected to Western blot analysis to assess the protein expression of bovine ACE2, NRP1, TMPRSS2, and bovine β-actin proteins. **(B)** Results of the qRT-PCR analysis of the bovine TMPRSS2 mRNA expression in the siRNA-Scrambled, siRNA-Furin, or the siRNA-BCoV-spike transfected MDBK cells. **(C)** Results of the Western blot band density of TMPRSS2 protein normalized to the β-actin. **(D)** Results of the qRT-PCR analysis of bovine ACE2 mRNA expression. **(E)** Western blot band density of ACE2 protein normalized with β-actin. **(F)** Results of the qRT-PCR analysis of bovine NRP1 mRNA expression. **(G)** Results of the Western blot band density of NRP1 protein normalized to the β-actin.

### 4. 9. The bta-miRNA-16 Overexpression enhances cytokines gene expressions in the context of BCoV infection

We investigated the impact of bta-miRNA-16a on the expression levels of the host cell cytokine production during BCoV infection. The MDBK cells were transfected with miR-16a, siRNA-Furin, and siRNA-spike, and the mRNA expression of IFN-α, IFN-β, IFN-γ, IL-6 and IL-10 were assessed by the qRT-PCR using the designed oligonucleotides listed in Table 2. Results showed significant up-regulation of IFN-α, IFN-β, and IFN-γ in the bta-miRNA-16a transfected group of cells compared to miRNA-Scr transfected cells (Fig 9A, 9B, 9C). The mRNA expression of host IL-6 and IL-10 was upregulated in the bta-miRNA-16a transfected group, particularly post-BCoV/Ent infection (Fig 9D, 9E). Meanwhile, the IFN-α, the IFN-β, and the IFN-γ expression levels were down-regulated in the siRNA-Furin and siRNA-spike transfected group of cells (Fig 9F, 9G, 9H). Although the expression levels of the IL-6 and IL-10 were upregulated in the siRNA-Furin and the siRNA-BCoV-S groups following BCoV/Ent infection, the overall expressions were lower compared to the siRNA-Scr transfected group of cells (Fig 9I, 9J).

**Fig 9:**
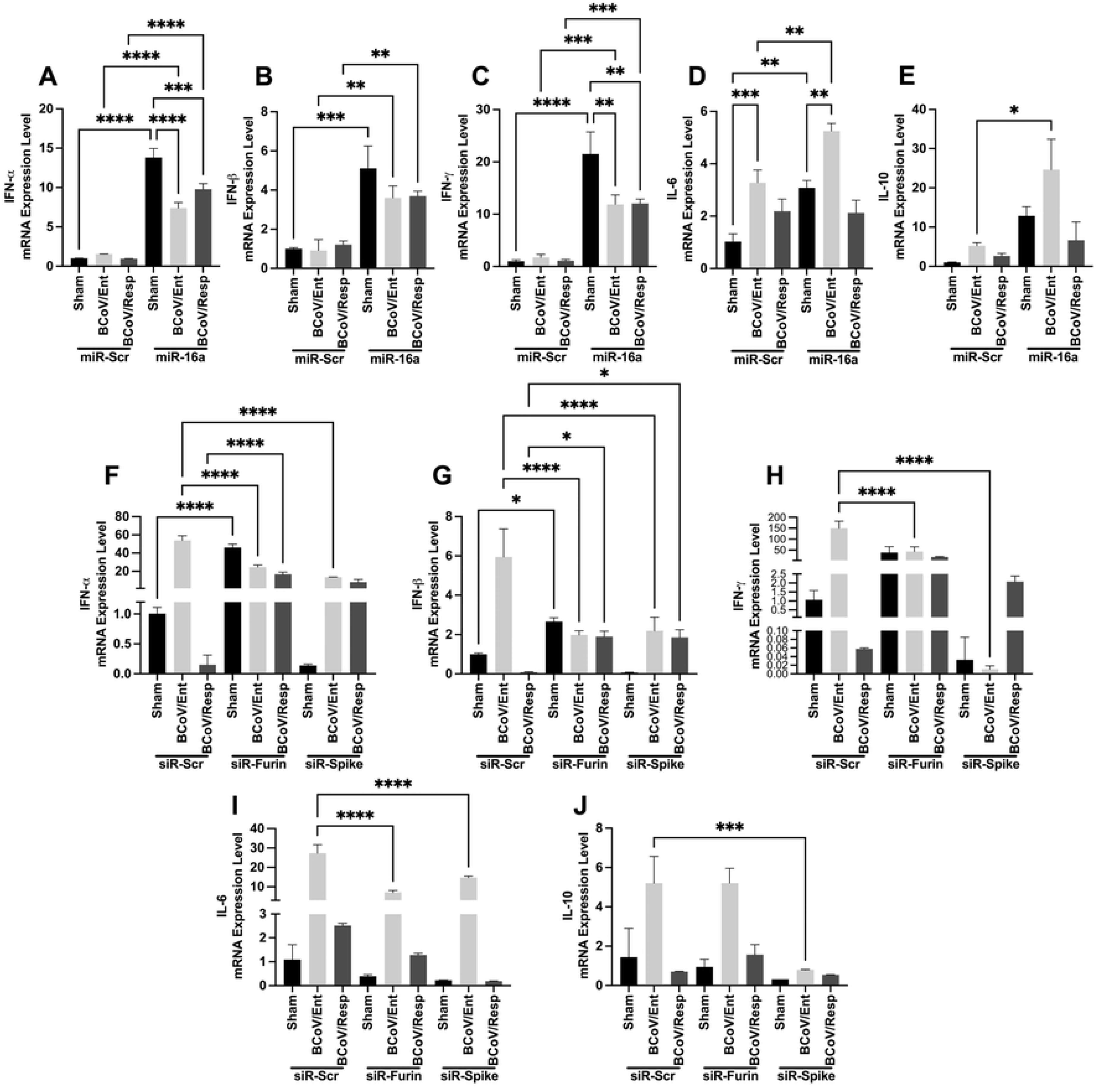
Activation of host cell cytokine-like Strome upon miR-16a overexpression during BCoV infection. The MDBK cells transfected with either the Scr-miRNA and bta-miRNA-16a, followed by BCoV/Ent or BCoV/Resp isolates infection. Results of the qRT-PCR analysis to quantify the mRNA expression of **(A)** IFN-α; **(B)** IFN-β; **(C)** IFN-γ; **(D)** IL-6; **(E)** and IL-10. **(F)** The MDBK cells transfected with Scr-siRNA control, siRNA-Furin, and siRNA-BCoV-Spike, followed by BCoV/Ent or BCoV/Resp isolates infection. Results of the qRT-PCR analysis to assess the mRNA expression levels of IFN-α; **(G)** IFN-β; **(H)** IFN-γ; **(I)** IL-6; **(J)** and IL-10.

## 4 Discussion

BCoV is one of the endemic viral pathogens in cattle populations worldwide, including the USA. (34). The virus has a dual respiratory/enteric tissue tropism (6). The mechanisms governing this dual tropism have not yet been well studied. The presence of viral receptors and other host cell proteases may partially explain this dual tropism; however, many aspects of the BCoV/host interaction, including their tissue tropism, have not yet been studied. However, the availability of viral receptors and some other host enzymes are not the sole agents that govern the success of viral replication or fully explain viral tropism. MicroRNAs are small RNA molecules that regulate the host genes at the post-translation levels. Previous studies showed that host miRNAs regulate the gene expressions of many host proteins during many viral infections, including SARS-CoV-2. Several host cell miRNAs, including miRNA-(155, 221, and 146a), could act as biological markers for SARS-CoV-2 infection; they also play essential roles in regulating the inflammatory responses during SARS-CoV-2 infection in humans (35, 36). However, the role of host cell miRNAs has not been extensively studied in the context of BCoV infection. The main aim of this study is to study the roles of some host miRNAs in BCoV replication, tissue tropism, and immune regulation/evasion. Our recent in silico prediction and bioinformatics search studies revealed that several host miRNAs could potentially play essential roles in the context of BCoV replication and explain at least in part the dual tropism of BCoV infection (37). In this study, we focused on the function of the beta-miRNA-16a that shows some differential expression profiles during the replication of either enteric or respiratory isolates of BCoV.

The miRNA/coronavirus interaction revealed some insights about the roles of these small RNA molecules in fine-tuning the virus replication and immune regulation. It was predicted that the hsa-miRNA-miR-8066 targeting SARS-CoV-2 nucleocapsid (N) protein leads to the inhibition of viral replication (38). Other studies showed at least seven host miRNAs (has-miRNA-(15a, 298, 497, 508, 1909, and 3130) targeting SARS-CoV-2-S glycoprotein at various locations, leading to substantial inhibition of the virus replication (39). The candidates of the host miRNA-16 family are involved in many cellular processes, including cell cycle arrest, tumor research, and immune regulation, particularly the TGFβ pathway, and they are also used as diagnostic and prognostic markers for cancer research (40, 41, 42, 43, 44, 45). Recent studies showed that the host miRNA-16-2-3p could be an excellent prognostic marker for SARS-CoV-2 infection in humans (46). However, the roles of bta-miRNA-16a in BCoV replication and immune regulation have never been studied yet.

Most coronaviruses, including members of Betacoronaviruses, use the host cell serine protease-2 (Furin) to cleave the spike glycoprotein at the PRRAR motif (47). This Furin/CoV/S targeting plays an important role in the CoV tissue tropism and molecular pathogenesis (48). It has been shown that the cleavage of SARS-CoV-2-S glycoprotein is an essential step in virus replication. The transmembrane serine protease 2 (TMPRSS2) proved to cleave SARS-CoV-2-S, and then another enzyme called Furin (another serin-protease-2) was responsible for the proteolysis of the Spike glycoprotein into the S1 and S2 domains (49). Those two simultaneous cleavage events are essential for most coronavirus replication, particularly in the entry of the virus into the host cells. To further confirm and consolidate this phenomenon, other researchers showed that deleting the Furin cleavage sites from the SARS-CoV-2-S substantially decreased the virus replication and impacted the downstream viral pathogenesis and tissue tropism (50, 51). Our data shows a differential expression of the bta-miRNA-16a in the BCoV/Ent and BCoV/Resp-infected MDBK and BEC cells (Fig. 2A-2D). There is a marked downregulation of the bta-miRNA-16a expression profile, especially in the case of the BCoV/Ent-infected cells. Thus, we believe the host responded to the BCoV/Ent infection by downregulating the bta-miRNA16a, which targets the 3’UTR of the host cell Furin. This bta-miRNA-16a targeting the Furin inhibits BCoV replication on the genome copy number level measured by qRT-PCR, on the viral protein’s levels including BCoV/S and BCoV/N protein, as well as on the virus infectivity levels as shown by plaque assay (Fig. 3D-3L). These data are very much consistent with SARS-CoV-2 loss of Furin cleavage sites that led to marked inhibition of SARS-CoV-2 replication (50).

Several approaches were used to validate some genes as targets for candidate miRNA, including bioinformatics search, western blot, and the dual luciferase assay on the potential gene 3’UTR construct carrying the seed region of the target miRNA. The comparison between the luciferase activity of the mutated 3’UTR construct of the miRNA target gene to that of the wild type construct carrying the seed region of the same miRNA candidate was used extensively as a benchmark for the miRNA/gene target validation (8, 28, 29). Our bioinformatic analysis for the bta-miRNA-16a shows the seed region of this miRNA is highly conserved among the Furin-3’UTR of several species of animals, including humans and bovine species (Fig. 4A). Our results show the Furin protein expression level is downregulated in the MDBK, and BEC transfected with bta-miRNA-16 compared to the scrambled miRNA transfected cells (Fig. 4C-4G). The dual luciferase assay results show marked inhibition of the luciferase activity in cells transfected with bta-miRNA-16a 3’UTR wild-type construct (Figure 4J). Thus, the Furin is considered a novel target for the bta-miRNA-16a.

Our data clearly show that infection with both BCoV/Ent and BCoV/Resp inhibits interferon (α, β, and λ) production (Fig. 9A-9C). However, the overexpression of the bta-miRNA-16a enhances cytokines gene expression (Fig. 9A-9C). Consistently, it was shown earlier that the BCoV-N protein inhibits the IFN-β production in the context of the RIG-I-like receptor (RLR) pathway (52). This confirms our findings about the dual actions of bta-miRNA-16a in the inhibition of BCoV infection and enhances the host immune response to counteract the viral infection.

One of the approaches to confirm the action of certain miRNA on their target genes is to use some siRNA to inhibit the same miRNA target region within these genes. To confirm the target validation of the bta-miRNA-16a on the host cell Furin and the BCoV-S glycoprotein, we designed two independent siRNA molecules to target the indicated regions. Our data shows that the application of siRNA-Furin and siRNA-S substantially inhibited BCoV replication in MDBK and BEC cells on the genome copy number, proteins (S and N) expression, and the virus infectivity by plaque assay (Fig. 5, 6). Our data is very much consistent with another research on other coronaviruses (CoVs), particularly SARS-CoV-2. Application of siRNAs targeting SARS-CoV-2-S glycoprotein significantly inhibited the virus replication (53). Similarly, using Furin inhibitors led to marked inhibition of SARS-CoV-2, which is considered a promising antiviral therapy for SARS-CoV-2 infection in humans (54, 55).

In addition to the host ACE2, several alternative host surface receptors have been suggested to promote coronavirus infection in host cells. The alternative receptors for ACE2 in coronavirus infection include the Neuropilin-1 (NRP1) (4, 56) and the CD147, a transmembrane glycoprotein expressed in epithelial and immune cells (57, 58). In most coronaviruses, including SARS-CoV-2, the spike protein undergoes two proteolytic cleavage steps after binding to ACE2. The first cleavage happened at the S1-S2 boundary, cleaved by host cell Furin. The virus entry still required a second cleavage by host cell proteases at the S2’ region, which is mostly done by TMPRSS2 (2). If the host cell has a low level of TMPRSS2 or if the virus-ACE2 complex does not meet TMPRSS2, the complex is taken into the cell by clathrin-mediated endocytosis, into the endolysosomes, where S2’s site is cleaved by cathepsins (59, 60). In this study, we showed that host Furin inhibition by bta-miR-16a or by siRNA-Furin down-regulates the ACE2 and NRP1 mRNA and protein expression (Fig. 7, 8). While no effect was observed on the TMPRSS2 expression at genomic or protein level (Fig. 7F, 7G, 8B, 8C). These findings suggest that bta-miR-16a down-regulates bovine host Furin, potentially inhibiting host ACE2 and NRP1 expression levels. Conversely, TMPRSS2 may compensate for these inhibitory effects of the bta-miRNA-16a on Furin through an alternative regulatory mechanism to rescue BCoV infection in host cells following Furin and ACE2 inhibition.

IL6 plays key roles in the regulation of some CoV replication, particularly SARS-CoV-2, and in the modulation of its immune response (61). SARS-CoV-2 infection triggers robust IL6 production, which induces the Cytokine storm (62). In a similar pattern, BCoV infection, particularly the BCoV/Ent isolate, induces upregulation of the IL6 expression levels (Fig. 9D). Furthermore, bta-miNRA-16a treatment resulted in the elevation of the IL6 expression in cells transfected with bta-miRNA-16a and infected with the BCoV-Ent isolate compared to the Scr-miRNA transfected cells and infected with BCoV/Ent isolate (Fig. 9D). These Findings demonstrate that bta-miRNA-16a overexpression significantly activates the host cytokine response. Moreover, the prevalence of BCoV/Ent infection leads to elevated expression of host cytokines and interferons.

Previous studies showed that the miRNA-16 family is involved in cell cycle control, cell survival, and apoptosis pathways (43). This miRNA also controls the balance of cell survival and apoptosis. Our data shows that bta-miRNA-16a enhances cell survival in the transfected cells and is infected with either the BCoV/Ent or BCoV/Resp compared to the Scr-miRNA mimics (Fig. S2A-S2D). This data could also be supported by the marked activation of host cytokine gene expression, as shown in (section 4.4). Thus, the high stimulation of the cytokines and the limitation of the CPE observed in the bta-miRNA-16a transfected group of cells and infected with BCoV contributed to the marked inhibition of the virus replication. In alignment with these findings, cells treated with the siRNA-BCoV-S showed less CPE compared to same type of cells transfected with the miRNA-Scr mimics (Fig. S2A-S2D and S3 Fig).

In summary, bta-miRNA-16a plays dual actions in the restriction of BCoV replication through targeting the spike glycoprotein and the host cell Furin. This bta-miRNA-16a/viral/host gene targeting inhibited the BCoV replication. It also stimulates the production of some host cytokines which contributed substantially to the cell survival in the bta-miRNA-16 transfected cells compared to the Scr-miRNA transfected cells (Fig 10).

**Fig 10:**
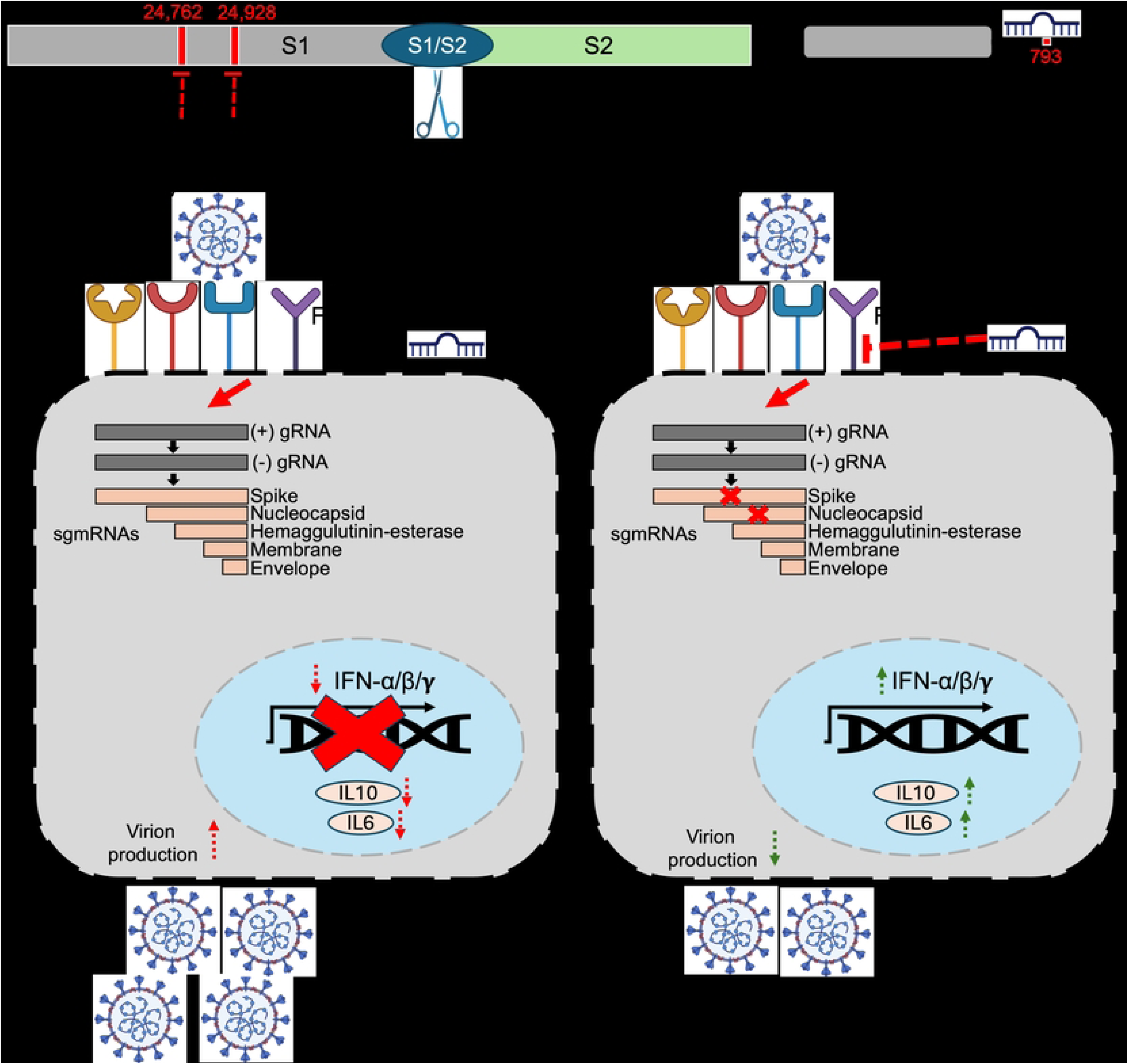
The proposed model of the mechanism of the dual actions of the host cell bta-miR-16a in restricting BCoV replication through targeting the BCoV-S glycoprotein and the host cell Furin and in the enhancement of cytokines expression. **(A)** Schematic representation of the BCoV-Spike protein, highlighting the S1 and S2 subunits and Furin cleavage site (S1/S2). The bta-miR-16a targets two regions within the S1 subunit, indicated in red (starting positions 24762 and 24928 respectively). **(B)** Schematic diagram of bovine Furin showing the bta-miRNA-16a target sites in the 3’UTR, red color indicated the starting nuclides of the seed region of the bta-miRNA-16a (starting position 793). **(C)** Illustration of bovine cells transfected with miRNA scrambled (miRNA-Scr) followed by BCoV infection. The BCoV viral genome released int the cytoplasm of the infected cells then the process of viral replication and the production of the nested sets of sg mRNAs initiated. BCoV infection resulted in marked inhibition in some host cytokines expression (IFN-α, IFN-β, IFN-γ, IL-6, and IL-10) but enhancing BCoV production. **(D)** Illustration of bovine cells transfected with the bta-miR-16a followed by infection with BCoV. The bta-miR-16a target host Furin at the cell surface reducing the BCoV activation through abolishing the spike glycoprotein cleavage. The overexpression of bta-miRNA-16 lead also to marked inhibition to the BCoV-S expression which ultimately reduced the production of other viral proteins particularly the BCoV-N. The bta-miRNA-16a targeting the BCoV-S and the host Furin resulted in marked inhibition on the production of new viral progenies. Meanwhile, this targeting resulted in marked increase in the expression of some host cell cytokines especially (IFN-α, IFN-β, IFN-γ, IL-6, and IL-10) which led to further inhibition of the viral progeny release.

Based on our findings in this study, we conclude that the bta-miRNA-16-a could be a promising diagnostic marker for BCoV infection. It also highlights the potential antiviral therapeutic potential of bta-miNRA-16a and its feasibility in designing some novel miRNA-based vaccines against BCoV in the future. Previously, it has been reported that miR-29a and miR-378b enhance the host DNA sensing Pathway in the presence of CpG motifs and can be used as an adjuvant for vaccines (63). Similarly, this study reveals that bta-miRNA-16a targets host Furin and enhances interferon production. Highlighting the importance of bta-miRNA-16a as a potential therapeutic avenue for BCoV infection in cattle.

## Supporting Information

**S1 Fig:**
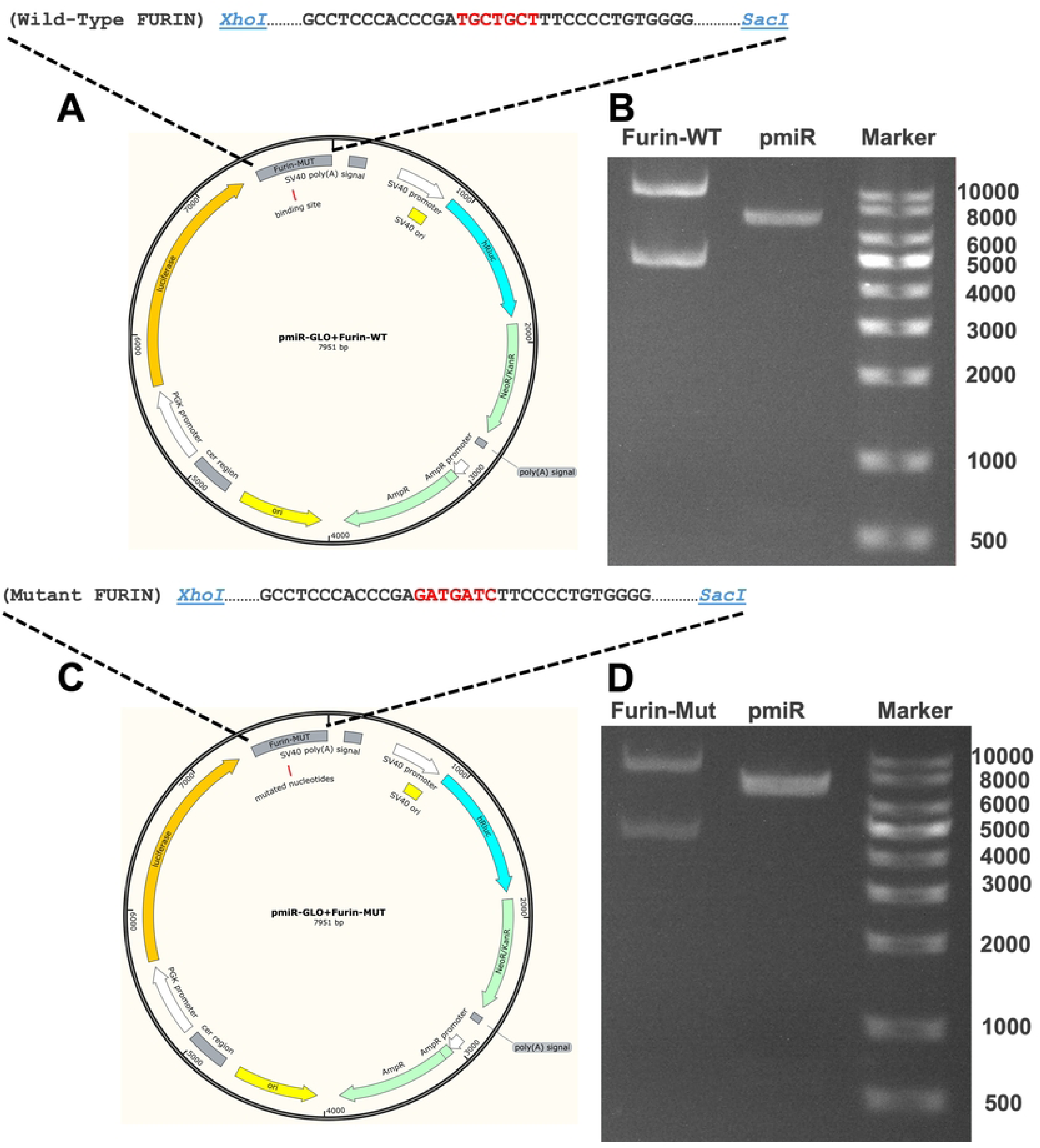
Design, construction, and confirmation of the pmiR-GLO dual Luciferase Reporter Plasmids containing either the wild type/mutated Furin-3’UTR. **(A)** Design of the pmiR-GLO plasmid with wild-type Furin inserted at *XhoI* and *SacI* restriction sites. **(B)** Confirmation of plasmid construct via restriction enzyme digestion by the agarose gel conformation. (Furin-WT: double digested with *XhoI* and *SacI*; empty pmiR-GLO vector; Marker). **(C)** Design of the pmiR-GLO plasmid with mutant Furin through the site-directed mutagenesis and insertion into the *XhoI* and *SacI* restriction sites of pmiR-GLO dual luciferase reporter vector. **(B)** Confirmation of mutant construct via agarose gel conformation. (Furin-Mut: double digested with *XhoI* and *SacI*; empty pmiR-GLO vector; Marker). About 200-1000ng plasmid was digested at 37°C for 30-60 minutes and analyzed through 1% agarose gel.

**S2 Fig:**
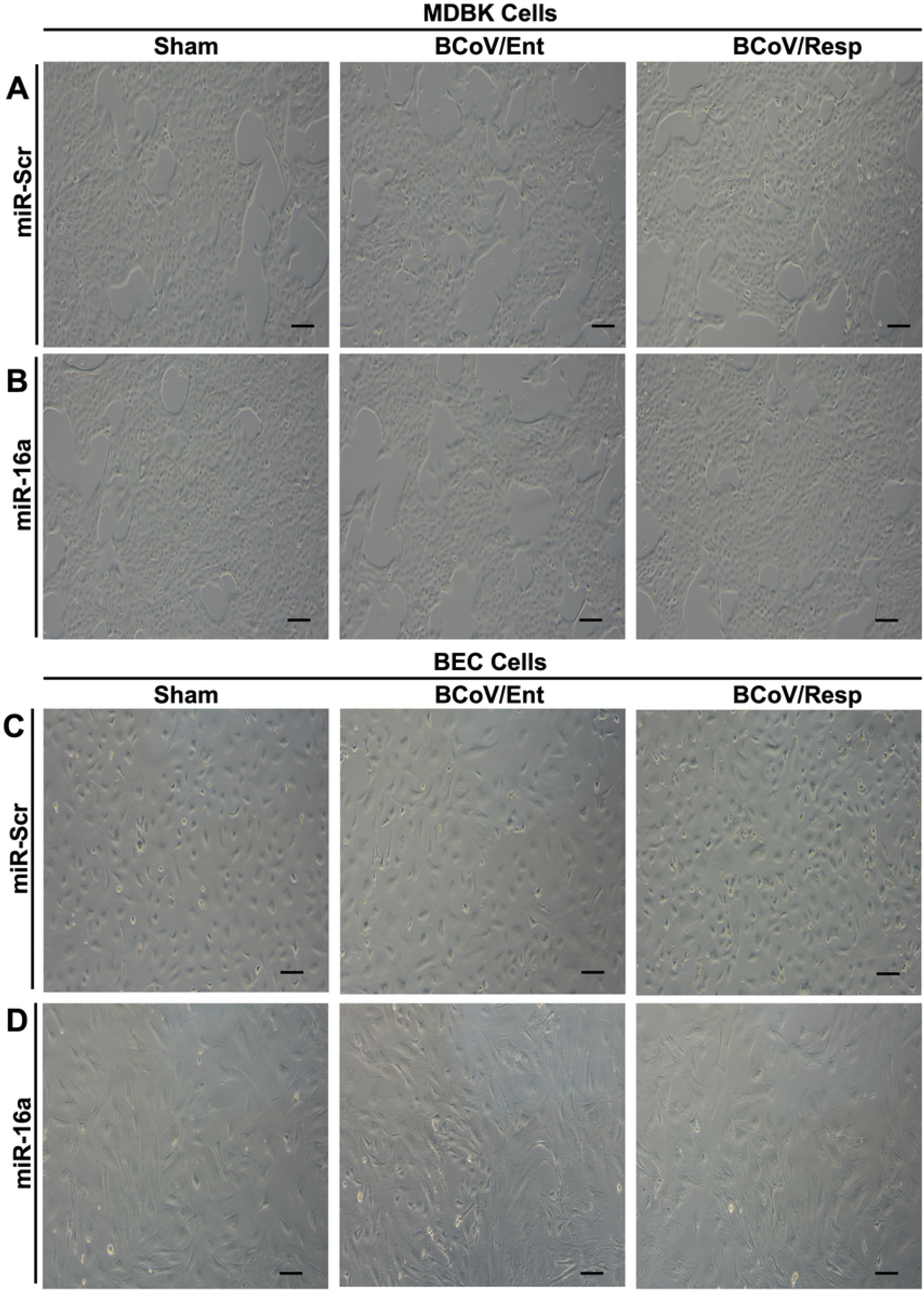
Morphology of bovine cells transfected with either the Sc-miRNA or the bta-miRNA-16a and then infected with either BCoV/Ent or BCoV/Resp isolates. **(A)** Results of the morphological examination of the MDBK cells transfected with the Scr miRNA as the relevant control and the bta-miRNA-16a, BCoV enteric (Ent), and BCoV respiratory (Resp) infected groups. **(B)** Results of the morphological examination of the MDBK cells transfected with the bta-miRNA-16a, BCoV/Ent, and BCoV/Resp infected groups. **(C)** Results of the morphological examination of the BEC cells transfected with the Scr-miRNA or the bta-miRNA-16a in the BCoV/Ent, and BCoV/Resp infected groups. **(D)** Results of the morphological examination of the BEC cells transfected with the bta-miR-16a or the Scr-miRNA control, BCoV/Ent, and BCoV/Resp infected groups. The bta-miR-16a was transfected for 24 hours, followed by 48-72 hours post-infection of the BCoV. All the images were captured at 10x magnification.

**S3 Fig:**
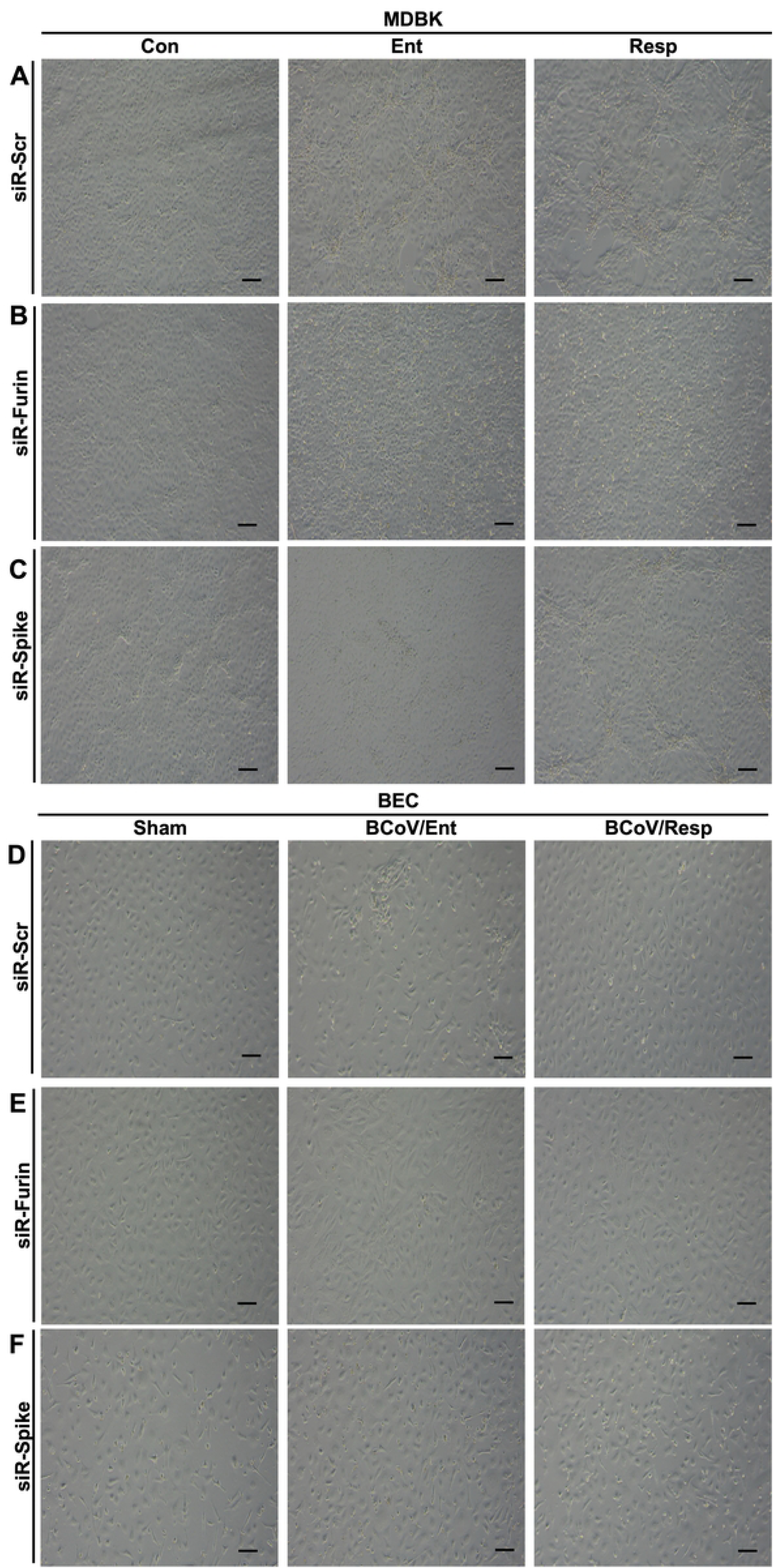
Morphology examination of the MDBK and the BEC Cells transfected with either the Scr-siRNA, the siRNA-Furin, or the siRNA-BCoV-S glycoprotein. **(A)** Morphological observation of the MDBK cells transfected with the Scr-siRNA in control (Sham), and infected with either the BCoV enteric (Ent), or the BCoV respiratory (Resp) isolates. **(B)** Morphological observation of the MDBK cells transfected with siRNA-Furin then infected with either BCoV/Ent or BCoV/Resp isolates. **(C)** Morphological observation of the the MDBK cells transfected with the siRNA-BCoV-Spike and then infected with either BCoV enteric, or BCoV respiratory infected groups. **(D)** Morphological observation of the BEC cells transfected with the Scr-siRNA, BCoV/Ent or BCoV/Resp isolates **(E)** Morphological observation of the BEC cells transfected with the siRNA-Furin and then infected with either the BCoV/Ent or BCoV/Resp isolates **(F)** Morphological observation of the BEC cells transfected with the siRNA-BCoV-Spike and then infected with either the BCoV/Ent or BCoV/Resp isolates. The siRNA-Furin and the siRNA-BCoV-Spike were transfected for 24 hours, followed by 48-72 hours post-infection of BCoV. All the images were taken at 10x magnification.

## Acknowledgments

We thank Drs. Udeni B. R. Balasuriya and Mariano Carossino, Louisiana State University, for kindly providing the Madine Darby Bovine Kidney (MDBK) and the Human rectal tumor -18 (HRT-18) cells used in this study. We also thank Dr. Aspen Workman from Animal Health Genomics Research Unit, USDA, A.R.S., U.S. Meat Animal Research, for kindly providing the Bovine Coronavirus Respiratory isolate used in this study.

## Author Contributions

M.G.H. conceptualized and designed the whole study, carried out some laboratory experiments, oversaw the entire research data analysis, wrote the manuscript, and submitted the manuscript. A.U.S. carried out the experiments and bioinformatic prediction, analyzed the data, and participated in writing the manuscript. All authors have read and agreed to the submitted version of the manuscript.

## Data availability statement

All data is available on request from the corresponding author.

